# Susceptible host availability modulates climate effects on dengue dynamics

**DOI:** 10.1101/2019.12.20.883363

**Authors:** Nicole Nova, Ethan R. Deyle, Marta S. Shocket, Andrew J. MacDonald, Marissa L. Childs, Martin Rypdal, George Sugihara, Erin A. Mordecai

## Abstract

Experiments and models suggest that climate affects mosquito-borne disease transmission. However, disease transmission involves complex nonlinear interactions between climate and population dynamics, which makes detecting climate drivers at the population level challenging. By analyzing incidence data, estimated susceptible population size, and climate data with methods based on nonlinear time series analysis (collectively referred to as empirical dynamic modeling), we identified drivers and their interactive effects on dengue dynamics in San Juan, Puerto Rico. Climatic forcing arose only when susceptible availability was high: temperature and rainfall had net positive and negative effects, respectively. By capturing mechanistic, nonlinear, and context-dependent effects of population susceptibility, temperature, and rainfall on dengue transmission empirically, our model improves forecast skill over recent, state-of-the-art models for dengue incidence. Together, these results provide empirical evidence that the interdependence of host population susceptibility and climate drive dengue dynamics in a nonlinear and complex, yet predictable way.

## INTRODUCTION

Ecological systems are complex, nonlinear, and dynamic. Understanding their mechanistic drivers is increasingly important in a rapidly changing world. Because of this complexity, mechanistic studies are often conducted in simplified, controlled environments at small scales. Connecting these experimental results with larger-scale emerging patterns remains a challenge but is nonetheless imperative for understanding the impacts of ecological change.

In concert with globalization and climate change, mosquito-borne diseases, and dengue in particular, are (re)emerging globally and spreading to higher latitudes (Kilpatrick & Randolph 2012; Ryan *et al.* 2019). Dengue virus—vectored primarily by urban *Aedes aegypti* (Kraemer *et al.* 2015)—places half of the global human population in 128 countries at risk of infection (Brady *et al.* 2012; Kraemer *et al.* 2019). In the absence of effective vaccines or treatments (Katzelnick *et al.* 2017a; Sridhar *et al.* 2018), public health agencies rely on vector control to reduce dengue transmission (Erlanger *et al.* 2008). Effective vector control interventions require understanding the mechanisms linking climate, vector ecology, disease transmission, and host population susceptibility to better predict disease outbreaks—a major challenge.

Since *Aedes* spp. mosquitoes are sensitive to climate, including temperature and rainfall (Stewart Ibarra *et al.* 2013; Mordecai *et al.* 2019), we expect temperature and rainfall to be important drivers of dengue outbreaks. Although temperature affects mosquito and viral traits in laboratory experiments (Watts *et al.* 1987; Lambrechts *et al.* 2011; Mordecai *et al.* 2017), the relationship between temperature and dengue incidence in the field has been ambiguous (Caldwell *et al.* 2020). Thus, temperature-dependent models have had mixed success predicting the timing and magnitudes of epidemics (Hii *et al.* 2012; Johansson *et al.* 2016; Johnson *et al.* 2018). Similarly, the rainfall–dengue relationship is complex, because the effect of rainfall on mosquitoes depends on local breeding habitat and human behavior. In some settings, rainfall fills container-breeding habitats for mosquitoes, increasing mosquito abundance and dengue incidence (Stewart Ibarra *et al.* 2013). By contrast, low rainfall can also facilitate dengue transmission by promoting water storage that serves as standing-water habitat for mosquitoes (Oliveira-lima *et al.* 2000). Further, heavy rainfall can reduce mosquito abundance by flushing out larvae (Koenraadt & Harrington 2008). The net effect of climate on dengue therefore depends on many different mechanisms and is highly context-dependent.

Disease incidence also depends nonlinearly on susceptible availability, because epidemic growth slows as the population of susceptible individuals is exhausted (Anderson & May 1979; Dushoff *et al.* 2004; Mina *et al.* 2015; Pitzer *et al.* 2015; Rypdal & Sugihara 2019). Further, susceptible availability may influence the effects of climate on dengue dynamics. However, such interactive effects are difficult to detect since susceptibility is difficult to observe, especially in endemic settings where multiple serotypes circulate and create a complex landscape of time-dependent and serotype-dependent immunity (Katzelnick *et al.* 2017b). Specifically, four serotypes of dengue regularly circulate in many regions: each provides long-term serotype-specific (homologous) immunity and short-term (heterologous) cross-protection against other serotypes (dos Santos *et al.* 2017; Jiménez-Silva *et al.* 2018; Hamel *et al.* 2019). Following a brief period of cross-protection, antibodies at a mid-range of titers can cause antibody-dependent enhancement of disease following heterologous, secondary infection, until titers decay to the point of nearly full heterologous susceptibility (Katzelnick *et al.* 2017b). Given this complex and dynamic immune landscape, directly detecting population susceptibility to circulating dengue virus at any point in time is difficult without longitudinal serology studies, which are not widely available (Gordon *et al.* 2013; Katzelnick *et al.* 2017b).

Previous prediction models of dengue outbreaks used phenomenological (Johansson *et al.* 2009b; Hii *et al.* 2012; Johnson *et al.* 2018) and mechanistic equation-based approaches (Tran *et al.* 2013; Liu-Helmersson *et al.* 2014; Morin *et al.* 2015; Mordecai *et al.* 2017), which may not fully capture interdependence between climate and susceptible availability. Phenomenological models may underperform when extrapolating past observed contexts, and equation-based mechanistic models rely on parameter estimates from laboratory studies engineered to isolate single mechanisms producing separate relationships between drivers and outcome, eliminating the complex interdependence at the population level. While laboratory studies provide robust validation of mechanisms (Lambrechts *et al.* 2011), the fixed relationships obtained from them do not necessarily translate into robust causal understanding for the complexity of field systems (Sugihara *et al.* 2012). Even if causality exists between two variables in such a system, their correlation can switch signs during different time periods, resulting in a net correlation of zero (Deyle *et al.* 2016b). This temporal variation in the direction of correlation results from the nonlinear, state-dependent relationship between the variables (i.e., the effect depends on another variable’s state). Conversely, even if two variables are consistently correlated, the association could be spurious due to a confounder.

To overcome these challenges, we used empirical dynamic modeling (EDM) (Sugihara *et al.* 2012)—a mechanistic, equation-free, data-driven approach that accounts for the context-dependence of ecological drivers to identify and model mechanisms driving dengue epidemics. EDM is based on reconstructing the essential system dynamics evident in time series, without assuming fixed relationships. This means relationships among variables can change through time to reflect that interactions among variables depend on context and system state. EDM does not require assumptions about the functional form of the model, but instead derives dynamic relationships empirically by constructing an attractor—a geometric object (i.e., curve or manifold) that embodies the rules for how relationships among variables change with respect to each other through time depending on system state (specific location on the attractor)—from time-series observations. Like a set of equations, the attractor encompasses the dynamics of a system, and thus can provide a mechanistic understanding of the system that is derived empirically, without requiring an *a priori* assumed set of equations.

Here, we used EDM and a proxy for susceptible population size (Rypdal & Sugihara 2019) to answer three questions: (1) Do temperature, rainfall, and/or inferred susceptible availability drive population-level dengue incidence? (2) Can we predict dengue dynamics using temperature and rainfall data and inferred susceptible availability? (3) What is the functional form of each climate–dengue relationship at the population level, and how is this relationship influenced by susceptible availability?

## METHODS

### Time series data

We obtained time series of weekly observations of dengue incidence (total number of new cases of all serotypes), average temperature (°C), and total rainfall (mm) in San Juan, Puerto Rico, for 19 seasons (1990/1991–2008/2009) spanning calendar week 18, 1990 to week 17, 2009 (Figure 1a–c) from the National Oceanic and Atmospheric Administration in November 2016 (http://dengueforecasting.noaa.gov/). We obtained data for four additional seasons (2009/2010–2012/2013) from Johnson *et al.* (2018) in April 2020 (https://github.com/lrjohnson0/vbdcast). Although dengue incidence data were also available for Iquitos, Peru (Johansson *et al.* 2019), we chose to focus on San Juan because the time series was longer, and therefore more amenable to EDM analyses (Munch *et al.* 2020).

**Figure 1.**
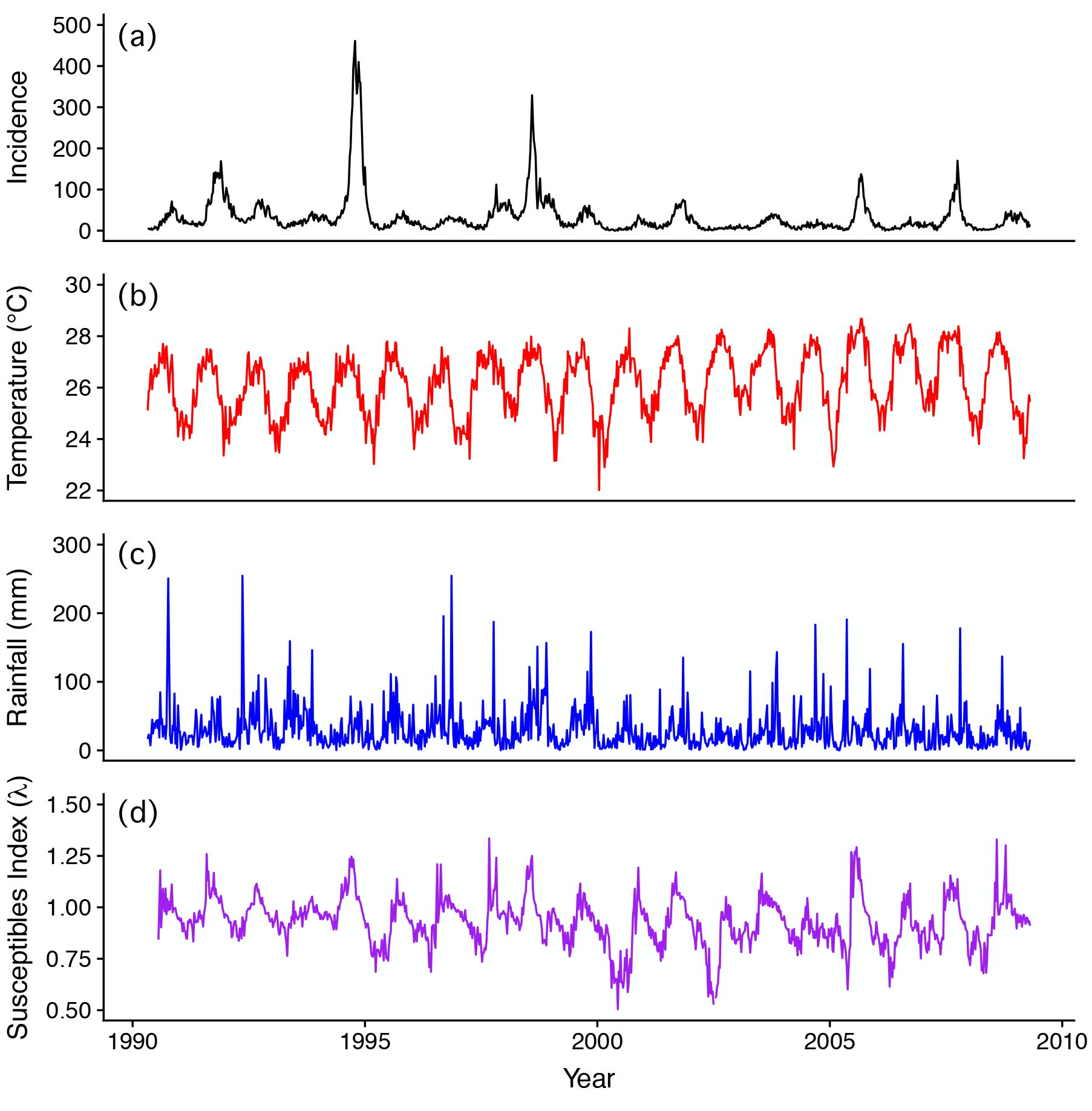
Dengue incidence, climate, and susceptibles index data. Time series (seasons 1990/1991–2008/2009) of (a) weekly dengue incidence (i.e., total number of cases per week), (b) weekly average temperature, (c) total weekly rainfall, and (d) a proxy for susceptible population size (see Supporting Information for details) in San Juan, Puerto Rico.

Direct measurements of susceptible availability are not available, so from weekly incidence data *I*(*t*), we estimated time-dependent growth rates: *λ* = *I*(*t* + Δ*t*)/ *I*(*t*). The growth rate, *λ*, is proportional to the effective reproduction number, *R*_eff_, and equivalent to *R*_eff_ if Δ*t* equals the average time between primary and secondary host infections. Vector-borne disease models show that *R*_eff_ is proportional to the geometric mean of the susceptible host population and the susceptible vector population: 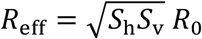, where *R*_0_ is the basic reproduction number (Zhao *et al.* 2020). Hence, 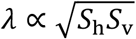 and *λ* can be used as a proxy for the susceptible population size at least during inter-outbreak periods where the transmission rate and *R*_0_ can be assumed to vary very little (Rypdal & Sugihara 2019).

We estimated *λ* by linear regression using the model *I*(*t* + Δ*t*) = *λ I*(*t*) for 12 time points in a 12-week running window (Δ*t* = 1 week). The model is robust to the window size (Rypdal & Sugihara 2019). In the discrete case, when *λ* < 1 the system is stable (inter-outbreak period) and when *λ* ≥ 1 then the system is unstable (outbreak period) (Supporting Information). We treated the resulting time series of *λ*, hereafter “susceptibles index” (Figure 1d), as a proxy for the susceptible population size when *λ* < 1, and a proxy for the combined effects of susceptible availability and *R*_0_ when *λ* ≥ 1.

### Empirical dynamic modeling (EDM)

EDM infers a system’s mechanistic underpinnings and predicts its dynamics using time series data of one or more variables to construct an attractor in state space (Figure S1). This procedure is called univariate (using lagged versions of a single variable time series) or multivariate state-space reconstruction (SSR). Properties of the attractor are assessed to examine characteristics of the system (Deyle & Sugihara 2011). We normalized each time series to zero mean and unit variance to remove measurement unit bias, ensuring the variables would be comparable and the attractor would not be distorted. All analyses were conducted in R version 3.5.1 (R Development Core Team 2018) and all EDM analyses were performed using package rEDM (Park *et al.* 2020).

To infer mechanisms, EDM should be applied in systems where there is evidence of underlying low-dimensional deterministic dynamics (Cummins *et al.* 2015). EDM assumptions are met when stochasticity is present (e.g., due to measurement noise, stochastic drivers, or unexplained variability) (Cenci *et al.* 2019; Munch *et al.* 2020), but the system cannot be entirely stochastic. To test for low-dimensional deterministic dynamics we performed univariate SSR for each variable, and used *simplex projection* (Sugihara & May 1990)—a type of nearest neighbor regression performed on an attractor—to check whether the system is forecastable beyond the skill of an autoregressive model—an indicator of underlying deterministic dynamics (Figures S2a and S4; Supporting Information). To test for nonlinear state dependence of a variable—the motivation behind EDM—we used the *S-map* test for nonlinearity (Sugihara 1994) (Figures S2b, c and S5; Supporting Information).

### EDM—Convergent cross-mapping

We used an EDM approach called *convergent cross-mapping* (CCM) (Sugihara *et al.* 2012) to identify drivers of dengue incidence. If two variables are causally related, then a multivariate attractor—where each variable in the system represents a dimension that traces the dynamics of the system—can be reconstructed (up to a practical limit) using lagged versions of just one of the variables (Figure S1). Based on Takens’ Theorem, this univariate “shadow attractor” preserves the structural and dynamic properties of the original multivariate attractor (Takens 1981; Sugihara *et al.* 2012). The concept behind CCM is that if temperature causes dengue incidence, then information about past temperature will be embedded in the dynamics of dengue, such that the shadow attractor produced using only incidence data allows us to accurately reconstruct temperature in the past. However, the converse scenario would not be true: since dengue does not cause temperature, the shadow attractor constructed using temperature data should not contain information to accurately reconstruct past dengue incidence (Supporting Information).

The critical criterion for estimating causal (directional) associations between two variables using CCM is checking that the cross-mapping skill (i.e., Pearson’s correlation coefficient, *ρ*, between predicted driver values using the univariate SSR of the response variable, and the observed driver values) monotonically increases and plateaus (i.e., converges) with the length of the response variable time series used in cross-mapping. We used the Kendall’s τ test as a significance test for convergence of cross-mapping skill using the Kendall package (McLeod 2011). If *τ* > 0 then there is convergence (Grziwotz *et al.* 2018).

We performed pairwise cross-correlations on the time series to investigate time-lagged relationships between potential drivers (i.e., temperature, rainfall, and susceptibles index) and dengue incidence using the tseries package (Trapletti & Hornik 2018). Based on these analyses (Figure S6), we applied a 9-week time lag between temperature and incidence, an averaged lag of 3–9 weeks for rainfall (i.e., the average rainfall over the preceding 3–9 weeks) to resemble standing water as mosquito breeding habitat over a longer time period, and a 5-week lag for the susceptibles index. These lags are proxies for the time delays of potential cause-and-effects and are consistent with results from field studies (Chen *et al.* 2010; Stewart Ibarra *et al.* 2013).

We assessed the strength of evidence for effects of potential drivers on dengue by comparing the CCM performance using the data with the performance of two null models that control for the seasonal trend (i.e., interannual mean) observed in all variables (Figure 2). These null models address the sensitivity of CCM to periodic fluctuations (i.e., seasonality), which can make two variables appear to be causally linked when instead they are simply synchronized by a seasonal confounder (Cobey & Baskerville 2016; Deyle *et al.* 2016a). In the first “seasonal” null model, we preserved the seasonal signal, but randomized the interannual anomalies (Deyle *et al.* 2016a). In the second, more conservative “Ebisuzaki” null model, we conserved any periodicity (beyond seasonal) and randomized the phases of Fourier-transformed time series (Ebisuzaki 1997). We tested for statistically significant differences in cross-mapping skill between the model that used the data *versus* the null models by performing Kolmogorov-Smirnov (K-S) tests after convergence.

**Figure 2.**
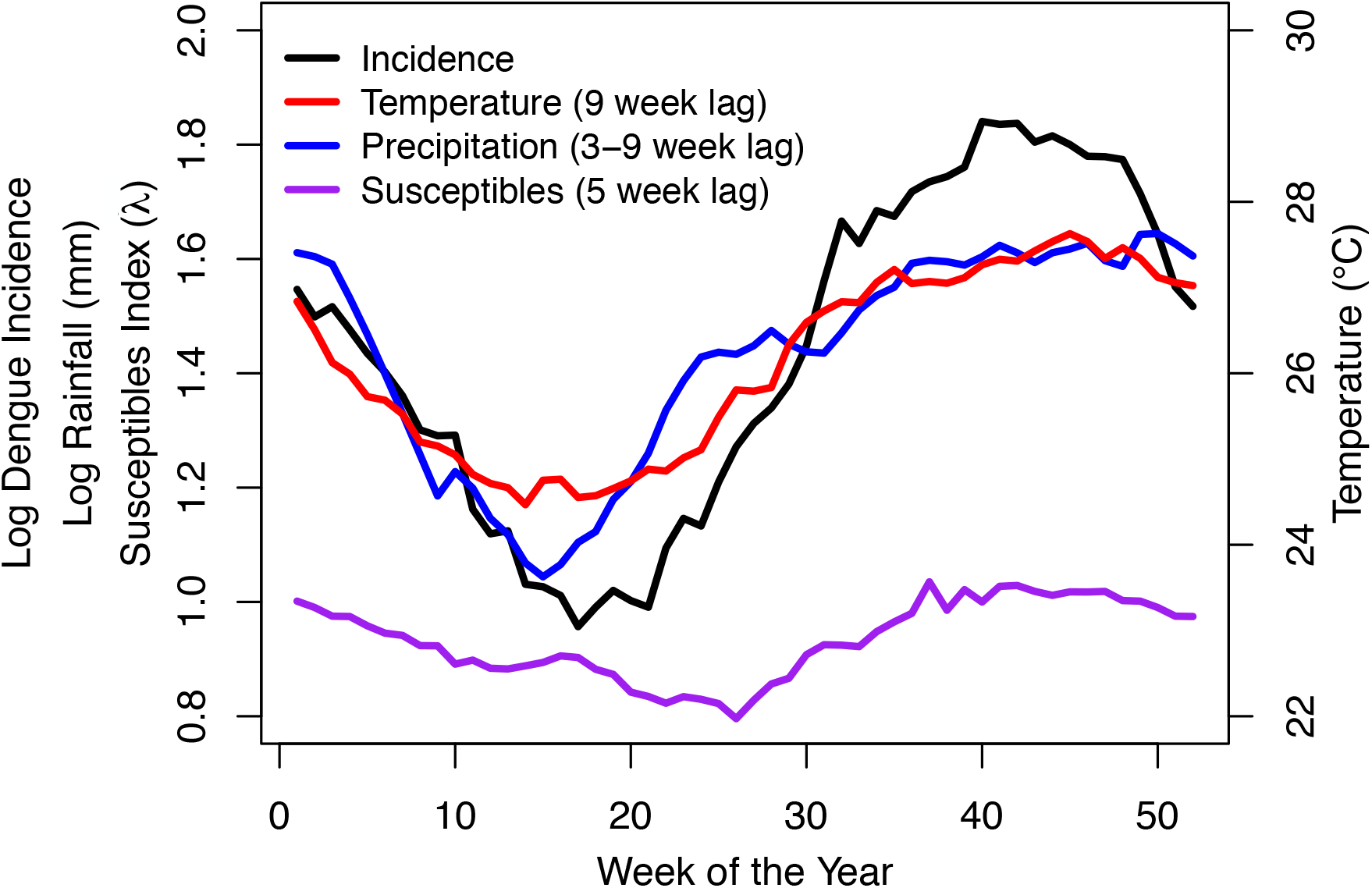
Seasonal trends and lags of dengue incidence and its drivers. The strong seasonal signal of dengue cases and other variables suggests potential causal lags between dengue incidence and temperature, rainfall, or the proxy for the susceptible population size. The lines represent interannual averages for each week of the year (i.e., calendar week) of dengue incidence (black), temperature shifted 9 weeks forward in time (red), average rainfall over the preceding 3–9 weeks and shifted 3 weeks forward in time (blue), and susceptibles index shifted 5 weeks forward in time (purple).

We also repeated CCM in the nonsensical, reverse-causal direction (e.g., to test whether incidence drives climate) as a control for potential spurious relationships generated by non-causal covariation (e.g., due to seasonality). This addresses the issue of synchrony, in which CCM can indicate bidirectional causality when one direction is false or nonsensical (Baskerville & Cobey 2017; Sugihara *et al.* 2017).

### EDM—Forecast improvement

We examined the predictive power of the drivers on dengue incidence by assessing how well we can predict dengue dynamics using temperature, rainfall, susceptibles index, and their combined effects. We used a combination of univariate SSR (i.e., with incidence data) and multivariate SSR to build forecasting models and to determine the improvement of forecasting using simplex projection when including different combinations of drivers (Dixon *et al.* 1999; Deyle *et al.* 2013, 2016a) (Supporting Information). We built the SSR forecasting models/attractors using the 1990/1991–2008/2009 season data (Figure 1) and made forecasts 8 weeks ahead. We assessed model forecasting performance using leave-one-out cross-validation.

Next, we evaluated out-of-sample forecasting performance of these models using testing data from four additional seasons (2009/2010–2012/2013). Predictions made on week zero for the first forecast of the 2009/2010–2012/2013 period (8 weeks ahead) came only from SSR using the 1990/1991–2008/2009 data. All subsequent weekly forecasts (8 weeks ahead) were made from updated SSR using all previous data, including past observations from the testing dataset.

Forecast uncertainty was evaluated by taking the density and morphology of the attractor into account. The more compact a simplex was and the less its starting position on the attractor mattered for the simplex projection, the more certain we were about our point estimate. Forecast variance was obtained from a distribution of weighted nearest neighbor regression from edges of simplexes constructed at various starting positions in the past.

Finally, we compared our top model performance with performance of previous models from 16 teams that participated in a dengue forecasting challenge (Johansson *et al.* 2019) and had access to the same data. To make a fair comparison, we followed the procedure as directed in the challenge (Supporting Information).

### EDM—Scenario exploration

In nonlinear systems, drivers generally have an effect that is state-dependent: the strength and direction of the effect depends on the current state of the system. Scenario exploration with multivariate EDM allowed us to assess the effect of a small change in temperature or rainfall on dengue incidence, across different states of the system. The outcome of these small changes allowed us to deduce the relationship between each climate driver and dengue incidence and how they depend on the system state. For each time step *t* we used S-maps (Sugihara 1994; Deyle *et al.* 2016a) to predict dengue incidence using a small increase (+Δ*X*) and a small decrease (−Δ*X*) of the observed value of driver *X*(*t*) (temperature or rainfall). For each putative climate driver, the difference in dengue predictions between these small changes is 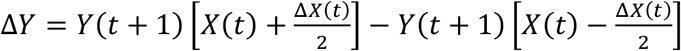, where *Y*(*t* + 1) is a function of *X* and all other state variables, and we used Δ*Y*/Δ*X* to approximate the effect of driver *X* at time *t*. We repeated this over all time steps in our time series for both temperature and rainfall to recover their approximate relationships with dengue incidence at different states of the system. Scenario exploration analyses were repeated across several model parameterizations to address potential sensitivity to parameter settings (Supporting Information).

## RESULTS

### Drivers of dengue dynamics

EDM showed that temperature, rainfall, and the susceptibles index drive dengue incidence since the convergence criterion was met (Kendall’s *τ* > 0, *P* < 0.01) in all three cases (Figure 3a–c). Rainfall and susceptibles index were significant drivers of dengue incidence beyond seasonality, as their effects were distinguishable from seasonal and Ebisuzaki null models (Figures 3b–c and S8b–c; K-S *P* < 0.0001). This implies statistically significant effects of both rainfall and the susceptibles index on dengue, which are not obscured by a periodic confounder. However, temperature was not a significant driver beyond seasonality (Figures 3a and S8a; K-S *P* = 0.90). We cannot rule out the possibility that the apparent forcing of temperature on dengue is due to a seasonal confounder. However, if no such confounder exists, then the seasonal trend in temperature, which accounts for most temperature variation in San Juan, drives the seasonal trend observed in dengue incidence. Compared to the other drivers, the converging cross-mapping skill of the temperature null models were relatively high (Figures 3 and S8), suggesting that temperature seasonality in each null model was a strong driver. Thus, seasonal temperature may be driving dengue dynamics, a result consistent with other studies (Huber *et al.* 2018; Robert *et al.* 2019).

**Figure 3.**
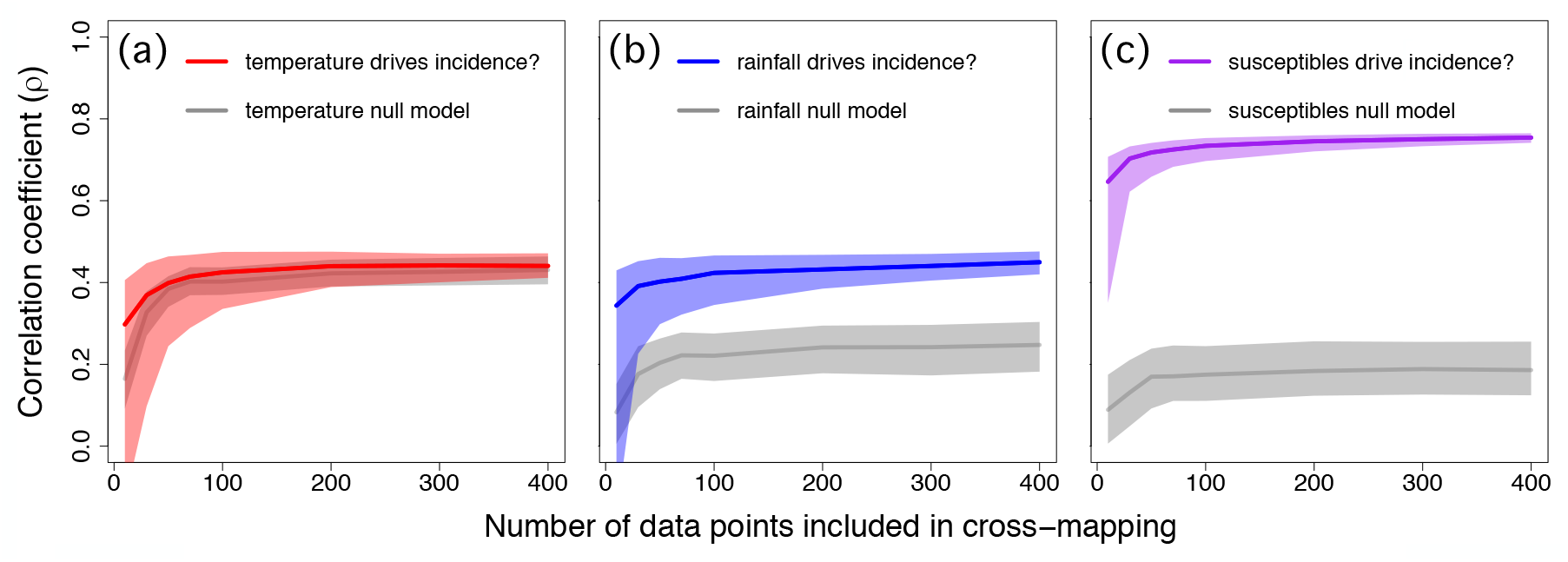
Climate and susceptibles index drive dengue incidence. Cross-mapping between dengue incidence and temperature (a; red), rainfall (b; blue), or susceptibles index (c; purple) display significant (Kendall’s *τ* > 0; *P* < 0.01) convergence in cross-mapping skill (i.e., *ρ* increases and reaches an asymptote) as the length of the time series increases (a signal of putative causality). Red, blue and purple shaded regions represent the 0.025 and 0.975 quantiles of bootstrapped time series segments. Grey shaded regions represent the 0.025 and 0.975 quantiles of the seasonal null distributions obtained from 500 runs of randomized time series with conserved seasonal trends (Deyle *et al.* 2016a). Solid lines represent medians of distributions. Rainfall and susceptibles index showed significant forcing above and beyond seasonal signal (K-S *P* < 0.0001), because cross-mapping of the true time series (blue and purple) are distinguishable from their respective null models (grey), whereas temperature forcing was not distinguishable from the null (K-S *P* = 0.90).

As expected, EDM tests for putative causality in the nonsensical directions— incidence driving temperature or rainfall—were not significant (i.e., no convergence; Figure S7, black lines). This result further supports the finding that temperature and rainfall drive dengue incidence, because their causal relationships were not confounded by spurious bidirectionality. The null models for the nonsensical directions of causality (Figure S7, grey lines) also displayed no convergence (completely flat), as expected (i.e., seasonality of dengue incidence does not drive seasonality of temperature or rainfall). However, seasonality (or any periodicity) of temperature, rainfall and susceptibles index drive dengue dynamics, shown by convergence of the seasonal and Ebisuzaki null models (grey lines in Figures 3 and S8).

### Predictive power of drivers

The multivariate SSR model using only temperature and rainfall data did not predict dengue incidence very well (*ρ* = 0.3839, RMSE = 47.72) although it captured the seasonality of the epidemics (Figure 4a). Forecasting skill doubled when the susceptibles index was included along with rainfall and temperature (*ρ* = 0.7547, RMSE = 37.40; Figure 4c), where timing and magnitude of epidemics were captured reasonably well. Dengue incidence prediction improved even further when incidence was added into the model with all drivers (*ρ* = 0.7662, RMSE = 37.14; Figure 4e). Dengue incidence was somewhat predictable using univariate SSR of incidence data alone (*ρ* = 0.4459, RMSE = 46.75; Figure 4g), suggesting that the dengue incidence time series contains information about its drivers, although limited. This points to some additional value of including the driver variables.

**Figure 4.**
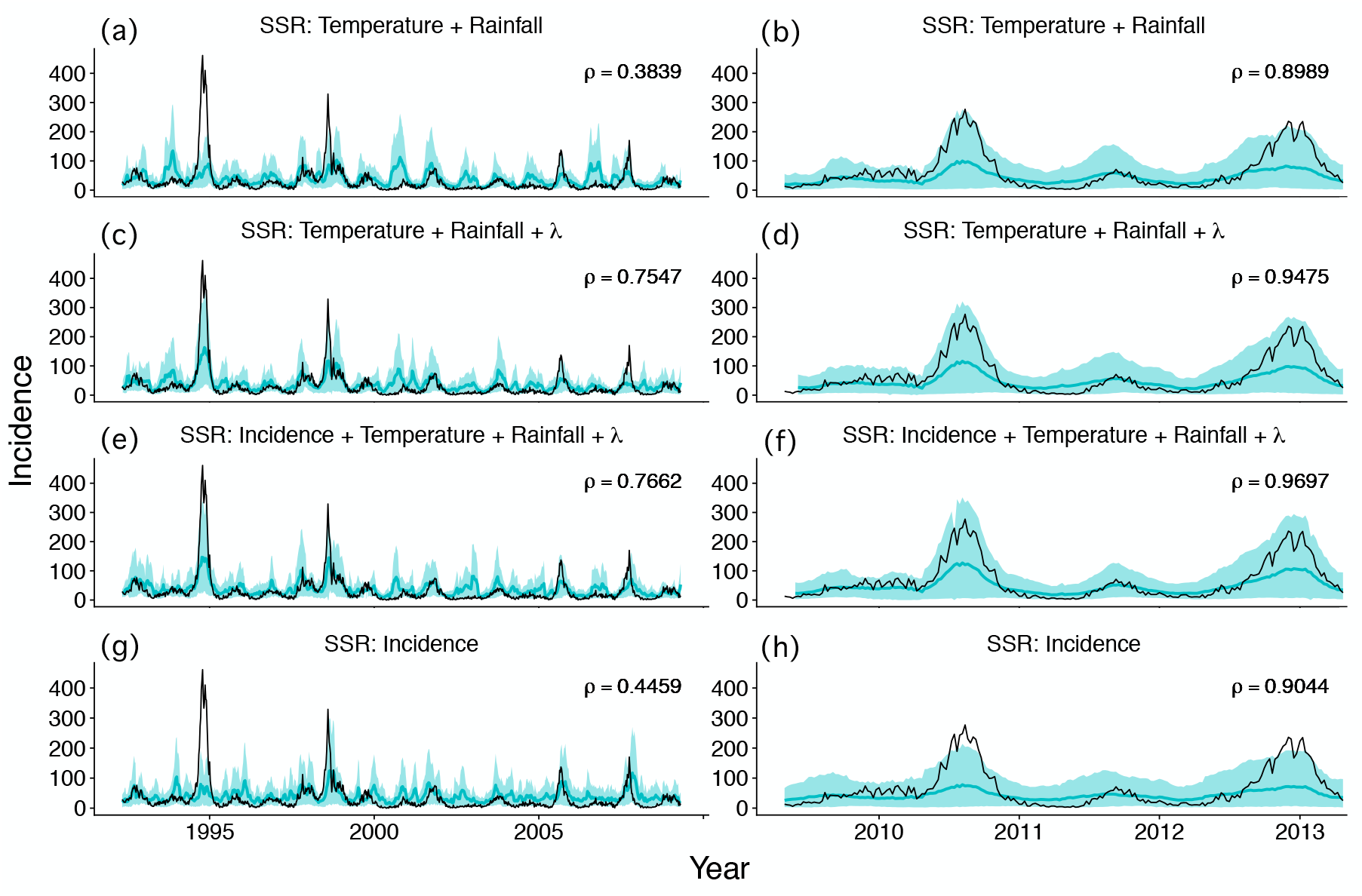
Predictive power of climate and susceptibles index (*λ*) on in-sample (left) and out-of-sample (right) dengue incidence. Forecasting results of incidence (8 weeks ahead) are shown in turquoise (solid lines represent the mean; shaded regions represent 90% confidence intervals) and observed incidence in black. (a, c, e, g) Time series for seasons 1990/1991–2008/2009 were used to construct SSR models for forecasts using leave-one-out cross-validation. (b, d, f, h) Data for seasons 2009/2010–2012/2013 were used to evaluate the SSR models constructed in a, c, e, and g, respectively, for out-of-sample forecasts. All SSR models (a–h) had significant forecasting skill (*ρ*) values (Fisher’s z-transformation *P* < 0.001).

We also evaluated the performance of the SSR models (Figure 4a, c, e, g) constructed using data from seasons 1990/1991–2008/2009 on external, testing data from 2009/2010–2012/2013 that were not used in SSR (Figure 4b, d, f, h). The average out-of-sample forecasting skill for each model for the testing seasons was higher than that of the 1990/1991–2008/2009 forecasts, although the errors were larger. The model using only temperature and rainfall displayed predictability (*ρ* = 0.8989, RMSE = 52.30; Figure 4b), the model that also included the susceptibles index improved predictions (*ρ* = 0.9475, RMSE = 52.12; Figure 4d), and the model that also included past incidence made highly accurate predictions (*ρ* = 0.9697, RMSE = 46.75; Figure 4f). The model that only included dengue incidence without the drivers was also predictive, although more error-prone (*ρ* = 0.9044, RMSE = 57.34; Figure 4h). All SSR models (Figure 4a–h) had significant forecasting skill (*ρ*) values (Fisher’s z-transformation *P* < 0.001).

The model with the highest prediction skill for the testing seasons (2009/2010–2012/2013), which included past climate, susceptibles index, and incidence data as predictors (Figure 4f), also outperformed models from the dengue forecasting challenge, including the ensemble model (Johansson *et al.* 2019) for predicting peak incidence, peak week, and seasonal incidence for all seasons on average (Tables S1–S2, Figures S9–S12). This demonstrates the benefit of the EDM approach for capturing the mechanistic, nonlinear, interdependent relationships among drivers over both equation-based mechanistic models and phenomenological models.

### State-dependent functional responses

As expected, we find state-dependent effects of temperature and rainfall with non-zero median effects. We found that temperature had a small positive median effect (2.88 cases/°C, Wilcox *P* < 0.001) on dengue incidence (Figure 5a). A positive effect is expected for the temperature range in Puerto Rico (Mordecai *et al.* 2017) (Figure 6e, black dashed lines), although the effect was occasionally much stronger, both positive and negative (Figure 5a, b). The large negative effects occurred only at the highest temperature values (as predicted by mechanistic models of temperature-dependent transmission), reinforced by a lower quantile regression with a strongly negative slope (Figure 5b, bottom dashed red line). However, positive effects occurred across the whole temperature range, which is limited to temperatures below the 29°C optimal temperature for transmission estimated from mathematical models and laboratory data (Mordecai *et al.* 2017).

**Figure 5.**
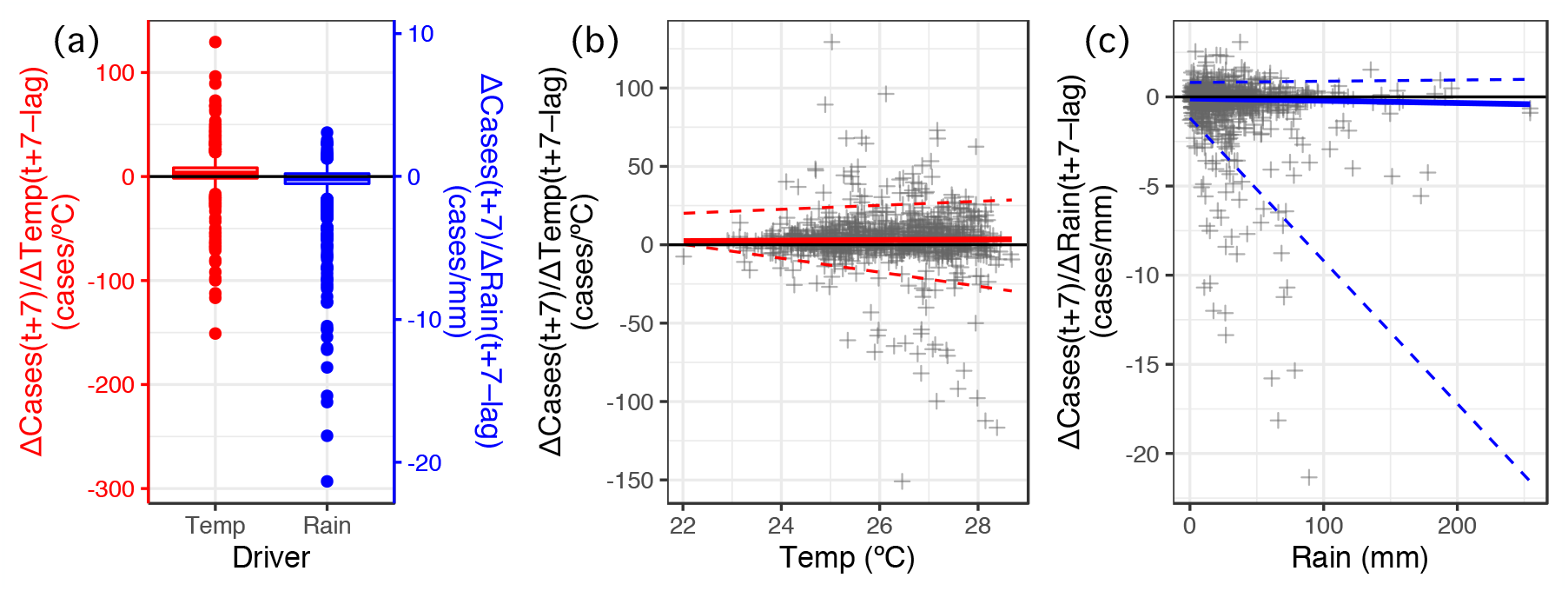
Temperature and rainfall show mixed effects on dengue incidence. Scenario exploration quantified the variable effect of changes in drivers on dengue. Boxplots show that the median effects of rainfall (Rain) and temperature (Temp) are small (close to zero), but drivers occasionally have strong impacts (a). To investigate climate driver functional responses, we plotted the rate of change of dengue incidence as a function of temperature (b) and rainfall (c). Red and blue lines represent regression on the median for temperature and rainfall, respectively, in a quantile regression. The dashed red and blue lines represent regression on the 0.05 and 0.95 quantiles of temperature and rainfall, respectively. Temperature has an overall positive effect on dengue incidence (median regression line of the rate of change is positive), but can also have large negative and positive effects (a, b). Rainfall has an overall negative effect (median regression line of the rate of change is negative), but can also have small positive and large negative effects (a, c).

**Figure 6.**
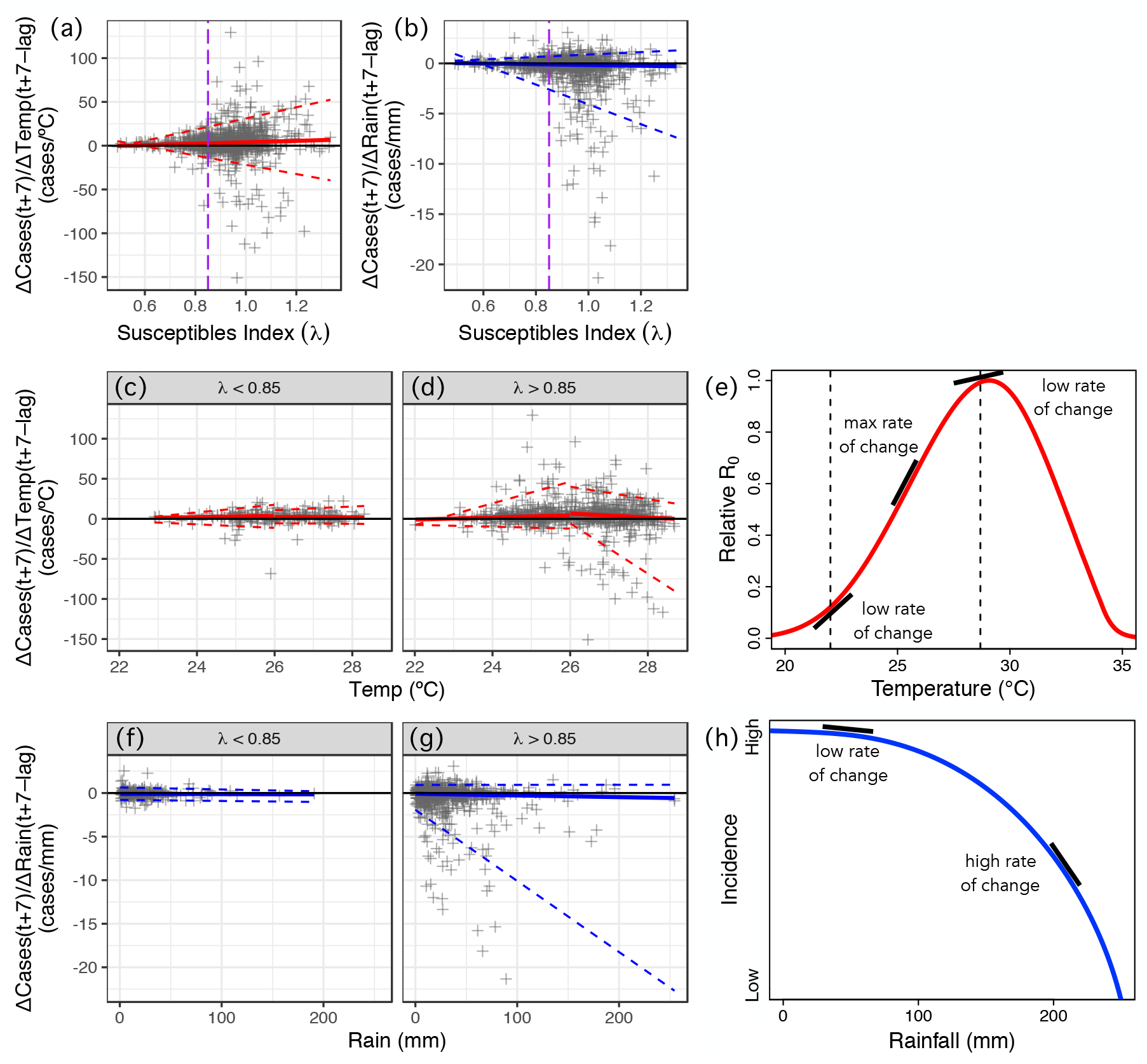
Temperature and rainfall effects on dengue incidence vary depending on the susceptible population size (*λ*). The effect of changes in temperature (a) and rainfall (b) against *λ* shows that driver effects are split around the threshold *λ* ≈ 0.85 (purple dashed line). The red and blue lines represent the median regression of temperature and rainfall effects, respectively, in a quantile regression (a–d, f, g). The dashed red and blue lines represent the 0.05 and 0.95 quantile regressions of temperature and rainfall effects, respectively (a–d, f, g). Neither driver has an effect on dengue incidence when susceptible availability is low (*λ* < 0.85; c, f). However, when *λ* > 0.85 climate effects are observed: temperature has mostly a positive effect (d), possibly sigmoidal in that temperature range (e), and rainfall has a negative effect (g), and conceptually a concave down functional response (h; black lines represent tangents, where the slope of the tangent is the rate of change). The effect of temperature on relative *R*_0_ of dengue assuming transmission via *Aedes aegypti* mosquitoes is unimodal (Mordecai *et al.* 2017) over a large temperature range (e; dashed lines indicate the minimum and maximum temperature values in the data of our study, black lines represent tangents, where the slope of the tangent is the rate of change of relative *R*_0_ of dengue as a function of temperature). Assuming that relative *R*_0_ is proportional to dengue incidence, our results suggest that the rate of change of dengue incidence is increasing until reaching a maximum and then decreasing (d; red median regression lines). However, even when driver effects are split at the evident threshold of *λ* = 0.85 (c, d, f, g), there are still many occurrences when the susceptible population size is sufficient large (*λ* > 0.85) but temperature and rainfall have no effect. In certain cases, temperature has even a negative effect on dengue (d).

Rainfall had a small negative median effect (−0.12 cases/mm, Wilcox *P* < 0.001), but occasionally had very large negative effects (Figure 5a, c). These large, negative effects of rainfall on dengue occurred when there was less than 100 mm of rain per week (Figure 5c), consistent with expectations that drought could lead to a high number of dengue cases due to water storage, which can provide mosquito breeding habitat (Oliveira-lima *et al.* 2000). There are also small positive effects of rainfall on dengue (Figure 5c), suggesting that overall the results showed competing effects of low to moderate rain providing standing water for mosquito breeding and humans storing water where mosquitoes can breed when there is drought or low rainfall.

These results suggest the strength and direction of the effects of climate on dengue dynamics depend on the state of the system. In addition to the nonlinear effects of climate drivers themselves on dengue incidence, another potential cause of state-dependent climate effects on dengue dynamics is the variation in the susceptible population size over time (Figure 6a, b). Outbreaks do not occur when there are too few susceptible people in the population. As expected, when the susceptibles index was small (*λ* < 0.85) incidence was insensitive to climate (Figure 6c, f). By contrast, when the susceptibles index was large (*λ* > 0.85), temperature and rainfall effects on dengue incidence appeared (Figure 6d, g). The gradual increase and decrease of the rate of change of dengue as a function of temperature (Figure 6d, red solid lines) aligned well with the changes in slope over the increasing part (Figure 6e, black dashed lines representing the temperature range in our study) of the unimodal temperature response curve for dengue transmission by *Ae. aegypti* developed previously (Mordecai *et al.* 2017). This is an important finding, since evidence of climate functional responses for disease dynamics is rare due to the difficulty of obtaining appropriately informative field data. It is possible that if we had temperature data ranging across a larger spectrum—possibly by assembling data across multiple climates—that the empirical functional response derived from EDM would also look unimodal. Further, when the susceptibles index was high, the slope of the relationship between rainfall and dengue incidence became more negative as rainfall increased, suggesting a concave-down effect of rainfall on incidence (Figure 6g, h). This relationship has been difficult to characterize in the field because of multiple, possibly context-dependent and lagged, mechanisms linking rainfall to dengue.

## DISCUSSION

High host susceptibility allows seasonal climate suitability to fuel large dengue epidemics in San Juan, Puerto Rico. The effects of climate and susceptibility are nonlinear, interdependent, and state-dependent, which makes inference from controlled mechanistic experiments, equation-based mechanistic models, or phenomenological models difficult. EDM provides an essential toolkit for identifying these drivers, quantifying their predictive power, and approximating their functional responses. In Puerto Rico, the causes of extensive interannual variability in dengue incidence have remained a mystery, despite hypotheses that climate and host susceptibility were involved. Here, we used EDM and a proxy for susceptible availability to disentangle nonlinear and interactive mechanisms driving disease dynamics.

We found that rainfall, susceptible availability, and plausibly temperature (via its seasonality) interact to drive dengue incidence. Combined, these three drivers predicted dengue incidence with high accuracy (Figure 4c). The EDM-based forecasting model outperformed 16 models and an ensemble model in a recently published dengue forecasting challenge (Johansson *et al.* 2019), suggesting that it could enhance dengue control efforts if surveillance efforts continue to report weekly case data. Finally, as expected from epidemiological theory, climate effects on dengue only appeared when susceptible availability exceeded a threshold (*λ* > 0.85; Figure 6).

The fact that climate effects are first observed when *λ* ≈ 0.85 (before the onset of an outbreak, *λ* = 1), suggests that rainfall, and possibly temperature, have an effect on the timing of an impending epidemic. Climate could drive the transmission rate, thus influencing *λ* (which is proportional to both susceptible population size and *R*_0_ when *λ* is close to 1), and therefore the timing of an outbreak could be attributed to the changes in transmission caused by seasonal climatic drivers (Rypdal & Sugihara 2019). The seasonality of temperature and rainfall had higher predictive skill than seasonality of susceptibles index (Figures 3 and S8, grey lines), further supporting that seasonality of incidence was associated with climate. However, the susceptibles index was critical for predicting dengue epidemic magnitudes (Figure 4c–f). Using the same data, Johnson *et al.* (2018) found that mechanistic models could predict the timing of seasonal epidemics, but that a phenomenological machine learning component was needed to capture interannual variation in epidemic magnitude. Our work suggests that the unobserved size of the susceptible population was a key missing link for predicting magnitude variation across years.

Previous studies have built models accounting for both susceptible availability and climate on dengue by reconstructing time series of susceptibles from a compartmental modeling framework (Metcalf *et al.* 2017). However, no previous studies on dengue have explored the interdependence between climate and susceptible population size. We showed that susceptible availability modifies climate effects on dengue: climate has negligible effects unless the susceptible population size is large enough (Figure 6). The interdependence of climate and population susceptibility has also been studied in diseases where the opposite effect was found. For example, climate effects on SARS-CoV-2 are expected to be negligible when susceptible availability is high in the early stage of the emerging pandemic (Baker *et al.* 2020). For influenza dynamics, population density in cities—potentially a proxy for susceptible availability—also modulated climate effects on disease transmission: climate effects were negligible in cities with high population densities (Dalziel *et al.* 2018).

Because dengue susceptibility is so complex—due to the serotype dynamics and time- and antibody titer-dependent cross-protection and enhancement (Katzelnick *et al.* 2017b)—total population density or size may not be a reasonable proxy for susceptible availability in dengue dynamics, and a direct mechanistic estimate of population susceptibility will likely never be widely available for most populations. Accordingly, it has been difficult for previous mechanistic models to capture susceptible dynamics for dengue and their interactions with climate. However, our approach provides a useful proxy that captures the susceptible population dynamics even in the absence of more detailed immunological information. By inferring the susceptibles index from incidence data, we were able to capture the strong influence of the susceptible availability on dengue dynamics, which in turn moderated the effect of climate on dengue dynamics. This result is expected from theory (Kermack & McKendrick 1927; Xu *et al.* 2017), but demonstrating it empirically is a unique contribution of this study.

Even when accounting for susceptible availability, the effects of temperature and rainfall on dengue were strongly state-dependent (Figure 6d, g). This result is potentially due to nonlinear effects of each climate driver (Figure 6e, h), interactions and correlations between temperature and rainfall, microclimate variation over space and time that is not captured by weekly averages, and complex lagged effects that are not captured by a single fixed lag (e.g., 9 weeks). In Puerto Rico, mosquitoes also breed in septic tanks year-round, allowing transmission to occur independently from rainfall (Mackay *et al.* 2009), thus weakening the rainfall–dengue negative relationship (Figure 6g). Some of this additional variation may be captured in the dengue incidence time series itself, which may explain why including it improves forecast skill over climate and susceptibility alone (Figure 4e, f). Despite this additional variation, our results are consistent with previous studies suggesting that dengue dynamics in Puerto Rico are positively associated with temperature (Johansson *et al.* 2009b; Barrera *et al.* 2011; Morin *et al.* 2015), and possibly negatively associated with rainfall (Johansson *et al.* 2009a; Morin *et al.* 2015), since most *Ae. aegypti* pupae in Puerto Rico are produced in human-made containers during periods of drought (Barrera *et al.* 2011).

The climate and incidence data used here have been used in multiple forecasting efforts, where ensemble approaches and approaches that incorporated mechanisms outperformed purely statistical approaches (Johansson *et al.* 2019). However, even the high-performing forecasting methods using the same dataset, and including (experimentally-derived) assumed mechanisms for the joint influence of climate and susceptibility on dengue dynamics, are still error-prone to the timing (on the order of weeks) and the magnitude (on the order of 50 cases) of intra-annual epidemics (Morin *et al.* 2015; Johansson *et al.* 2019). Mechanisms isolated independently in controlled experiments do not necessarily scale up to the population level, and susceptible dynamics derived from compartmental models may be too simple to properly capture true susceptibility at the population level for dengue (Katzelnick *et al.* 2017b). EDM allowed us to infer mechanisms empirically from population-level data, and accounted for the population-level interdependence between climate and susceptible availability for forecasting, which probably contributed to our model outperforming previous forecasting models and ensembles (Table S1).

Connecting climate and dengue at the population level is challenging, because causal relationships are likely to be nonlinear and state-dependent. A toolkit of methods for testing hypotheses, understanding mechanisms, and making predictions is essential for understanding disease dynamics in complex, natural populations. Ultimately, understanding how climate-driven vector-borne diseases are influenced by other variables, such as susceptible population size, is important for optimizing vector control under critical conditions where climate might spark epidemics. Our approach, using EDM and an inferred proxy for the susceptible population size from data, confirmed that climate has nonlinear, seasonal effects on dengue epidemics in San Juan, Puerto Rico, but only above a certain threshold of susceptible availability. The mechanisms inferred from EDM could be applied to understand and predict future ecological responses to changing environments, including dengue epidemics in a world undergoing rapid environmental change.

## ACKNOWLEDGEMENTS

We thank Giulio De Leo, Marcus Feldman, Dmitri Petrov, and members of the Fukami, Mordecai, Peay, and Sugihara labs for helpful feedback. NN was supported by the Bing Fellowship in Honor of Paul Ehrlich and the Stanford Data Science Scholars program. ERD and GS were supported by the National Science Foundation (NSF) DEB-1655203, NSF-ABI-1667584, DoD-Strategic Environmental Research and Development Program (SERDP) 15 RC-2509; Lenfest Foundation Award 00028335 and the McQuown Chair in Natural Sciences, University of California, San Diego. MSS and EAM were supported by an NSF Ecology and Evolution of Infectious Diseases grant (DEB-1518681). EAM was also supported by an NSF Rapid Response Research grant (RAPID 1640780), an NIH NIGMS R35 MIRA award (R35GM133439), the Stanford University Woods Institute for the Environment Environmental Ventures Program, the Hellman Faculty Fellowship, a Stanford King Center Seed Grant, and the Terman Fellowship. AJM was supported by an NSF Postdoctoral Research Fellowship in Biology (1611767). MLC was supported by the Lindsay Family E-IPER Fellowship and Illich-Sadowsky Interdisciplinary Graduate Fellowship.

