## Supporting Information for "Susceptible host availability modulates climate effects on dengue dynamics"

#### SUPPLEMENTARY METHODS

##### Derivation of a proxy for susceptible population size

Consider a simple Susceptible-Infected-Recovered (SIR) model (Kermack & McKendrick 1927),

$$\frac{dS}{dt} = -\beta SI \qquad (1)$$
$$\frac{dI}{dt} = (\beta S - \gamma)I$$
$$\frac{dR}{dt} = \gamma I$$

where $S$ denotes the number of susceptibles, $I$ the number of infected individuals, and $R$ the number of recovered individuals at a given time $t$, and $\beta$ is the transmission rate, and $\gamma$ is the recovery rate. If we assume that the total population size, $N$, is known and closed ($S + I + R = N$), then the vector ($S(t)$, $I(t)$) is sufficient to describe the time-varying state of the population.

In the more general case, we can further assume that $\boldsymbol{I} = (I_1, \ldots, I_n)$ and $\boldsymbol{S} = (S_1, \ldots, S_n)$ represent different infected and susceptible (human and/or vector) subpopulations, respectively. Thus, a generalization of Eq. 1 can be represented as: $(\dot{\boldsymbol{I}}, \dot{\boldsymbol{S}}) = \boldsymbol{F}(\boldsymbol{I}, \boldsymbol{S}; \beta)$, where $\boldsymbol{F}$ represents a nonlinear function. During an inter-outbreak period (i.e., a disease-free state) $\boldsymbol{I} = 0$ and we have that $\boldsymbol{F}(0, \boldsymbol{S}; \beta) = 0$, which represents a system equilibrium independent of $\boldsymbol{S}$ and $\beta$. The stability of a system is determined by the Jacobian matrix and its eigenvalues. The stability of an inter-outbreak period is thus determined by the Jacobian matrix $D\boldsymbol{F}(0, \boldsymbol{S}; \beta)$. Since $\boldsymbol{F}$ is constant when $\boldsymbol{I} = 0$, we have that $\frac{\partial \boldsymbol{F}}{\partial \boldsymbol{S}} = 0$ and thus the Jacobian matrix is singular (its determinant is zero) when $\boldsymbol{I} = 0$. Thus, we have that $\boldsymbol{I}$ can be linearized as $\dot{\boldsymbol{I}} = \frac{\partial \boldsymbol{F}}{\partial \boldsymbol{I}}(\boldsymbol{S}, \beta)\boldsymbol{I}$, and the stability of this system is determined by the leading eigenvalue $\lambda$ of $\frac{\partial \boldsymbol{F}}{\partial \boldsymbol{I}}$, which only depends on $\boldsymbol{S}$ and $\beta$. For example, in the case of the model described by Eq. 1, the leading eigenvalue would be:

$$\lambda = \beta S - \gamma \tag{2}$$

In the case of a vector-borne disease model, such as the Bailey−Dietz model (Bailey 1975; Dietz 1975), we have that during a disease-free period, when there are no infected human and vector populations ($I_h = I_v = 0$), we have that

$$\lambda = \frac{\gamma_h}{2} \pm \sqrt{\left(\frac{\gamma_h}{2}\right)^2 + \beta_1 \beta_2 S_h S_v} \tag{3}$$

where $\beta_1$ is the transmission rate of the pathogen from infected vectors to susceptible humans, and $\beta_2$ is the transmission rate of the pathogen from infected human to susceptible vector (Rypdal & Sugihara 2019).

In the continuous case, because $\lambda$ is an eigenvalue, when $\lambda < 0$ the system is stable (inter-outbreak period), and when $\lambda \geq 0$ the system is unstable (outbreak period). In the discrete case, however, we instead have that when $\lambda < 1$ the system is stable (inter-outbreak period) and when $\lambda \geq 1$ then the system is unstable (outbreak period).

Due to the linear relationship (Eq. 2, 3), $\lambda$ can act as a proxy for the proportion of the population that is susceptible over time in the inter-outbreak period (and hence it works even when the total population size changes over time), and can be inferred from incidence data alone, since from Eq. 1–3 it follows that $\frac{dI}{dt} = \lambda I$. For more details please see (Rypdal & Sugihara 2019). Although this proxy does include both vector and human susceptibility in the case of vector-borne diseases (Eq. 3), we are making the assumption that these two populations correlate positively, at least under long time scales, and so changes in $\lambda$ would correspond to changes in the human susceptible population size.

This approach has previously been used to infer year-to-year changes of susceptible availability during inter-outbreak periods (Rypdal & Sugihara 2019). Although few dengue cases occur during the inter-outbreak period, incidence during this time contains information about the susceptible population size in the next outbreak, and other factors that influence $R_{\text{eff}}$ can be assumed to vary very little over these periods. In this study, we used $\lambda$ as a proxy for the susceptible population size during both inter-outbreak and outbreak periods. Although we can assume that $\lambda$ captures changes in the susceptible population size during the inter-outbreak period (when $\lambda < 1$), it may also depend on other factors that influence $R_{\text{eff}}$ (i.e., transmission rate and climate) during the outbreak periods (when $\lambda \geq 1$). However, whenever $\lambda < 1$ the system is still in the inter-outbreak period and we can be more certain that the susceptibles index does represent changes in susceptible availability over time (results presented in Figure 6).

In the forecasting models (Figure 4) we used $\lambda$ from both the inter-outbreak and the outbreak periods, so $\lambda$ may not always have been proportional to the susceptible population size since it would have also depended on $R_0$. Nevertheless, adding the susceptibles index to our models improved our forecasting results of dengue incidence (Figure 4), and so even when $\lambda \geq 1$ the susceptibles index contains valuable predictable information of the system.

#### Empirical dynamic modeling (EDM)

##### EDM—Simplex projection

Simplex projection is a type of nearest neighbor regression that can be used to predict future states of a system on an attractor (i.e., a set of values in state space towards which a dynamical system tends to evolve) (Sugihara & May 1990). After the attractor is built using univariate or multivariate state-space reconstruction (SSR; Figure S1), simplex projection works as follows. For a given starting point, $t$, on the attractor, we use the nearest neighbors of that point to form a shape or a "simplex" with dimension $E + 1$, where $E$ denotes the number of "embedding dimensions" (i.e., the number of dimensions in state space, e.g., Figure S3). For example, a simplex could be a tetrahedron with 4 nearest neighbor points in a 3-dimensional state space (Figure S2a). We then project each nearest neighbor point forward one step in time to generate a simplex for the next time step, and the central point in that simplex is the predicted value for the next time step $t + 1$. The predicted value at $t + 1$ can be compared to the observed value at $t + 1$ from the raw time series, as a measure of the predictability of the constructed shadow attractor. In general, the longer the time series (i.e., more data points) the denser the shadow attractor and the smaller the simplex becomes, leading to higher predictability. This is called "convergence" because the prediction skill increases and plateaus at a maximum, with the length of the time series, and it is an essential criterion for finding a plausible causal relationship in convergent cross-mapping (see main text section EDM—Convergent cross-mapping).

**EDM—S-map test for nonlinearity**

S-map is a type of linear regression performed on an attractor to test if a system is nonlinear (Sugihara 1994). The concept behind S-map is that if local points are more predictive than non-local points, then that is an indicator of state dependence. S-map works as follows. First, a univariate attractor of a variable ($y$) is constructed using univariate SSR (Figure S1b). Second, S-map predictions of each data point are performed using a weighted multivariable linear regression model

$$y(t+1) = \beta_0 + \beta_1 y(t-E+1) + \cdots + \beta_E y(t)$$

where $t$ represents the time point for a particular data point used for prediction (and $t+1$ represents the time point for a particular data point to be predicted), $\beta_0 \ldots \beta_E$ represent the weights, and $E$ denotes the number of embedding dimensions in SSR (e.g., Figure S3). The weights are Jacobian coefficients computed using singular value decomposition and proportional to $e^{-\theta d/\bar{d}}$, where $d$ is the 2-

norm distance between the point to be predicted and each predictor point, and $\bar{d}$ is the mean of $d$. Thus, the weight depends on how close a predictor point is to the point to be predicted and the attractor localization parameter, $\theta$. This localization parameter, $\theta$, controls how much weight the local points have compared to all points in global state space. If $\theta = 0$ then all attractor points are weighted equally (linearity, no state dependence). However, if $\theta > 0$ then the local points are weighted more than nonlocal points (nonlinearity, state dependence). Thus, if the optimal $\theta > 0$ for a univariate SSR, then the variable is a part of a nonlinear dynamic system (i.e., location of a predictor point on the attractor matters; Figure S2b).

**EDM—Convergent cross-mapping**

Cross-correlation analyses alone do not provide enough evidence for causality between time-dependent variables. Instead, we used an EDM approach called convergent cross-mapping (CCM) (Sugihara *et al.* 2012). In this method, rather than correlating values in independent time series, we use the time series to reconstruct the dynamics of the system as a whole by building an attractor using SSR, to predict future states of the system, and to infer directional associations between variables.

Before diving into an intuitive explanation of CCM, imagine a multivariate SSR where each variable represents a dimension that traces the dynamics of the system through time in high-dimensional space (Figure S1c) (Sugihara *et al.* 2012). For instance, the variables dengue incidence, temperature, rainfall, and the susceptibles index ($\lambda$) form a 4-dimensional attractor using such a multivariate SSR approach (Figure S1d–f). Downstream multivariate SSR analyses can be used for many different purposes, such as scenario exploration (see EDM—Scenario exploration) and forecasting state-dependent interactions (Deyle *et al.* 2016b) to explore and forecast mechanistic interactions. Basically, the goal is to get a better mechanistic understanding of the system by including a response variable and its drivers in such a multivariate SSR framework. However, we usually do not know *a priori* what the drivers are; we only have hypothesized drivers, and we need to test for plausible causality before using them in the multivariate SSR explained above. Here, CCM is used to test the hypothesized causal drivers.

If two variables $X$ and $Y$ are causally related in a system, then the multivariate attractor can be reconstructed using lagged versions of just the response variable $Y$, i.e., $Y(t), Y(t-\tau), Y(t-2\tau), \ldots, Y(t-(E-1)\tau)$, where $\tau$ denotes the lag period for variable $Y$ (Figure S1b), and $E$ represents the number of embedding dimensions that maximizes the predictive power of the attractor (Figure S3). According to Takens' Theorem, this univariate "shadow attractor" preserves the structural and dynamic properties of the original multivariate attractor (Takens 1981). Here, we used simplex projection (Sugihara & May 1990) again to measure the predictive power of the attractor and to estimate $E$ (Figure S3). CCM detects causal relationships between variables $X$ and $Y$ as follows. First, a univariate shadow attractor is constructed using the response variable $Y$. Next, we try to predict a certain point on the attractor at time $t$ using $E + 1$ nearest neighbors of a point at a previous time step, e.g., $t - 1$, as in simplex projection. After locating the nearest neighbors and identifying their time points we look up the corresponding values of those same time points on the putative driver variable time series. We then take the average of those values, and obtain the predicted value of the driver $X$ at time $t$. This is then compared to the observed driver value at time $t$ to calculate the prediction skill, i.e., the Pearson correlation coefficient between predicted and observed values. If the prediction skill is non-zero and improves with the length of the time series used in CCM until reaching a plateau (indicating convergence), then there is support for a causal relationship between $X$ and $Y$. Relative to correlational approaches, this method has the advantage of being state-dependent and time-dependent, because it uses the full state space of the dynamic system (i.e., the values of each variable at each time point).

**EDM—Forecast improvement**

We used EDM forecast improvement to see how well temperature, rainfall, and the estimated susceptibles index can predict dengue incidence dynamics and to assess the strength of the putative causal relationships. Using multivariate SSR (Figure S1c), a shadow attractor is constructed from a combination of the driver variables and their lagged counterparts (e.g., $X_1(t),\ X_1(t-\tau),\ X_2(t), \ldots ,\ X_3(t)$ for driver variables $X_1$, $X_2$ and $X_3$ and time lag $\tau$) so that the total embedding dimension equals $E$ (i.e., the optimal embedding dimension for a univariate SSR of the response variable). We can then compare the prediction accuracy of dengue incidence of such a multivariate SSR using just the drivers with the prediction accuracy of the univariate SSR using just the response variable. This comparison in prediction skill will also inform us of the strength of the putative causal relationship.

We first constructed a shadow attractor using just lagged versions of the dengue incidence time series for up to $E$ dimensions (a univariate SSR) and used simplex projection to determine the attractor's prediction skill for dengue incidence, based on Pearson's correlation coefficient between predicted and observed values. We then constructed a shadow attractor using dengue incidence, lagged versions of itself, and temperature up to $E$ dimensions and determined the prediction skill. We repeated the process for rainfall alone, susceptibles index alone, and for all drivers combined, in each case comparing predictive skill with the univariate SSR using dengue incidence alone.

**EDM—Scenario exploration**

Due to the high auto-correlation in dengue cases from week to week, sequential time lags of 1 week contain very little new information and thus do not efficiently reconstruct the attractor (Casdagli *et al.* 1991). Therefore, we used a time-step of 7 weeks, where the auto-correlation function $ACF(\tau) = \rho(X(t),\ X(t+\tau))$ is approximately $1/\sqrt{e}$ (in practice this is the highest autocorrelation that allows us to distinguish the nonlinear predictability using simplex projection from autocorrelation; Figure S4). This ensures that each lag-coordinate used in the reconstruction contains unique information. Larger separation between lag coordinates ($\tau > 7$ weeks) could also work well in theory, but reduce the effective time series length. Additionally, we used a more conservative cross-validation scheme than the typical leave-one-out cross-validation approach: since nearby points in time have an extremely strong statistical relationship, we excluded all observations from within 6 months of target points for making predictions with scenario exploration.

Univariate analysis of dengue incidence time series revealed a large embedding $E$ (i.e., many lagged versions of dengue incidence are needed as dimensions to unfold the attractor dynamics). This is not ideal for scenario exploration analysis as the more lag coordinates used as proxy variables, the more opaque the meaning of the scenario exploration analysis becomes. To reduce $E$, we replaced the multiple lagged versions of dengue cases with total cases of dengue for the previous year (a moving one-year window average). We found the optimal multivariate embedding for scenario exploration analysis by taking the identified drivers—temperature, rainfall, and susceptibles index—together with incidence (cases per week), incidence lagged by 7 weeks, and the total cases of dengue for the previous year (mentioned above). When including total yearly dengue cases for the previous year, we were able to represent the system with fewer dimensions, since the total yearly dengue cases collapsed the embedding by half without losing predictability. This is because the total yearly dengue cases dimension represented the remaining half of the total number of dimensions of lagged cases by 7 weeks. Finally, we performed standard S-map analysis to identify the optimal $\theta$ for predicting future weekly cases.

#### Dengue forecasting challenge

We followed the procedure as instructed in the Dengue Forecasting Project (https://dengueforecasting.noaa.gov/). All details are specified in (Johansson *et al.* 2019). In brief, we made seven seasonal forecasts starting at season weeks $n$ = {0, 4, 8, ..., 24} for all four training seasons (2005/2006–2008/2009) based on past data (1990/1991–2004/2005) (Figures S10 and S12, Table S2), and four testing seasons (2009/2010–2012/2013) based on past data (1990/1991–2008/2009) (Figures S9 and S11, Table S1).

At each season week, we made forecasts (weekly predictions) on the attractor built using past data. The forecasting window ranged from 1 week ahead up to the last season week, 52. We computed a logarithmic score based on model performance, accounting for prediction variability, by having the model output be the probability that a prediction outcome falls in a particular bin for each forecast target per season: peak week (binned by season week 1–52), peak incidence (binned by 50 cases and last bin was 500 or more), and total number of cases in a season (seasonal incidence, binned by 1000 cases and last bin was 10,000 or more). The logarithmic score per season was the average logarithm of the probability assigned to the observed outcome bin, $p_i$, for all seven forecasts in set $n$: $S = \frac{1}{7}\sum_{i=1}^{7}\log(p_i)$.

Let $\mathbf{f}_n$ be the vector of the weekly future predictions and past observed data for the seasonal forecasting performed each week in set $n$. Several forecasts were made by performing simplex projection on the attractor built from data up to week $n$. To obtain a distribution of forecasts $\mathbf{f_n}$ we varied the site of simplex construction and performed the nearest neighbor regression step on the edges of the simplexes (as described in the main text). Based on the distribution of $\mathbf{f_n}$ the probabilities $p_i$ from our model predictions were computed as follows for each target:

1. <u>Peak week:</u> For each $\mathbf{f_n}$ we noted the week of peak incidence, and repeated this for all $\mathbf{f_n}$ to obtain a distribution of predicted peak weeks. The resulting empirical distribution of peak week was approximated as a skewed normal distribution. From the distribution density function we noted the probability of the actual observed peak week, $p_i$.
2. <u>Peak incidence:</u> For all forecasts, we obtained the maximum of all entries of each $\mathbf{f_n}$ to obtain a distribution of the predicted peak incidence. The resulting empirical distribution of peak incidence was approximated as a skewed normal distribution. The distribution probability density was split into intervals as the specified bins, and the integral of each interval turned into probabilities $\mathbf{p}$. From $\mathbf{p}$ we selected $p_i$ based on the bin that contained the observed peak incidence.
3. <u>Seasonal incidence:</u> For all forecasts, we summed all entries of each $\mathbf{f_n}$ to obtain a distribution of the predicted total number of cases for that season. The resulting empirical distribution of seasonal incidence was approximated as a skewed normal distribution and its density was split into intervals as the specified bins, and the integral of each interval turned into probabilities $\mathbf{p}$.

From **p** we selected $p_i$ based on the bin that contained the observed seasonal incidence.

We then computed the average logarithmic score for all four training seasons (2005/2006–2008/2009), and the average logarithmic score for all four testing seasons (2009/2010–2012/2013) and those scores are shown in Tables S2 and S1, respectively.

### SUPPLEMENTARY FIGURES

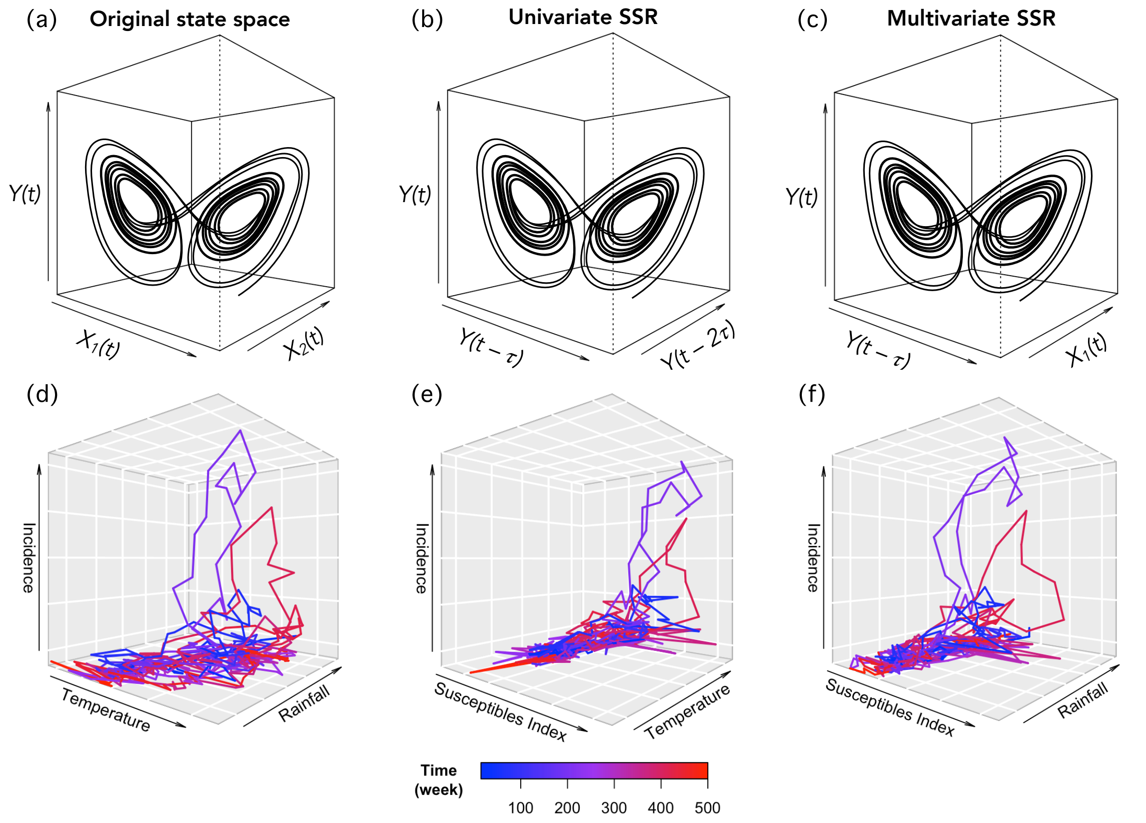


**Figure S1. Variations of state-space reconstruction (SSR).** The system's trajectory through state space can be plotted in three dimensions, for example, with response variable $Y(t)$ and driver variables $X_1(t)$ and $X_2(t)$ (a). A similar trajectory can be plotted with univariate SSR coordinates; $Y(t), Y(t-\tau)$, and $Y(t-2\tau)$, where lags of $Y$ replace driver variables $X_1(t)$ and $X_2(t)$ (b). The trajectory can also be plotted in multivariate SSR coordinates, e.g., in terms of $Y(t)$ (and lagged version of it) and $X_1(t)$ (c). The univariate SSR can be compared to the multivariate SSR to determine the magnitude of the forecasting improvement when including drivers (Deyle *et al.* 2013). To provide an example, we plotted the attractor of a multivariate

SSR of our system using the four variables: incidence (dengue cases per week), temperature (weekly average), rainfall (weekly total), and susceptibles index (proxy for susceptible population size), where the times series have been standardized (d–f).

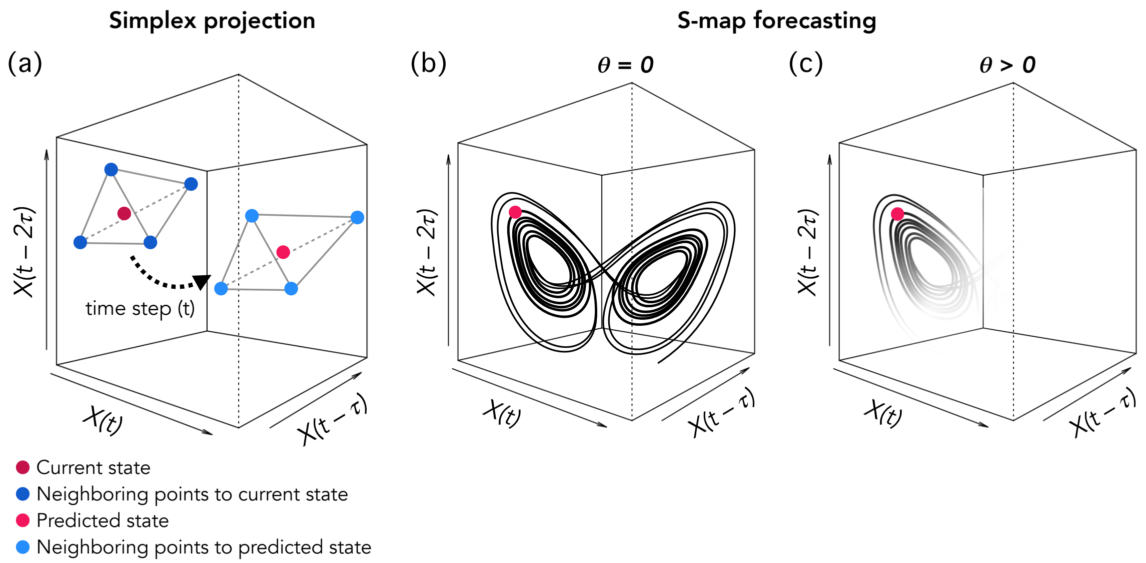


**Figure S2. Simplex projection and S-map forecasting.** (a) Simplex projection is a forecasting method for state-space reconstruction (SSR). The embedding dimension ($E$) in this example is three. Simplex projection uses a library of past states to select $E$+1 nearest neighbors to the current state of the system. The resulting simplex (a tetrahedron in this example) is then projected forward in time (time step $t$) to predict the future state based on the temporal changes of the neighboring states gathered from the observed history of the system time series data. (b, c) S-map is another forecasting method for SSR. It can be used to test if a system is nonlinear. If local points on the attractor (close to the point to be predicted) are more predictive than points further away, then the system is state-dependent and nonlinear (c). The localization parameter, $\theta$, controls how much weight the local points have compared to all other points on the attractor: (b) If $\theta = 0$ then all points are weighted equally (system is linear), (c) but if $\theta > 0$ then the local points are weighted more than other points (system is nonlinear).

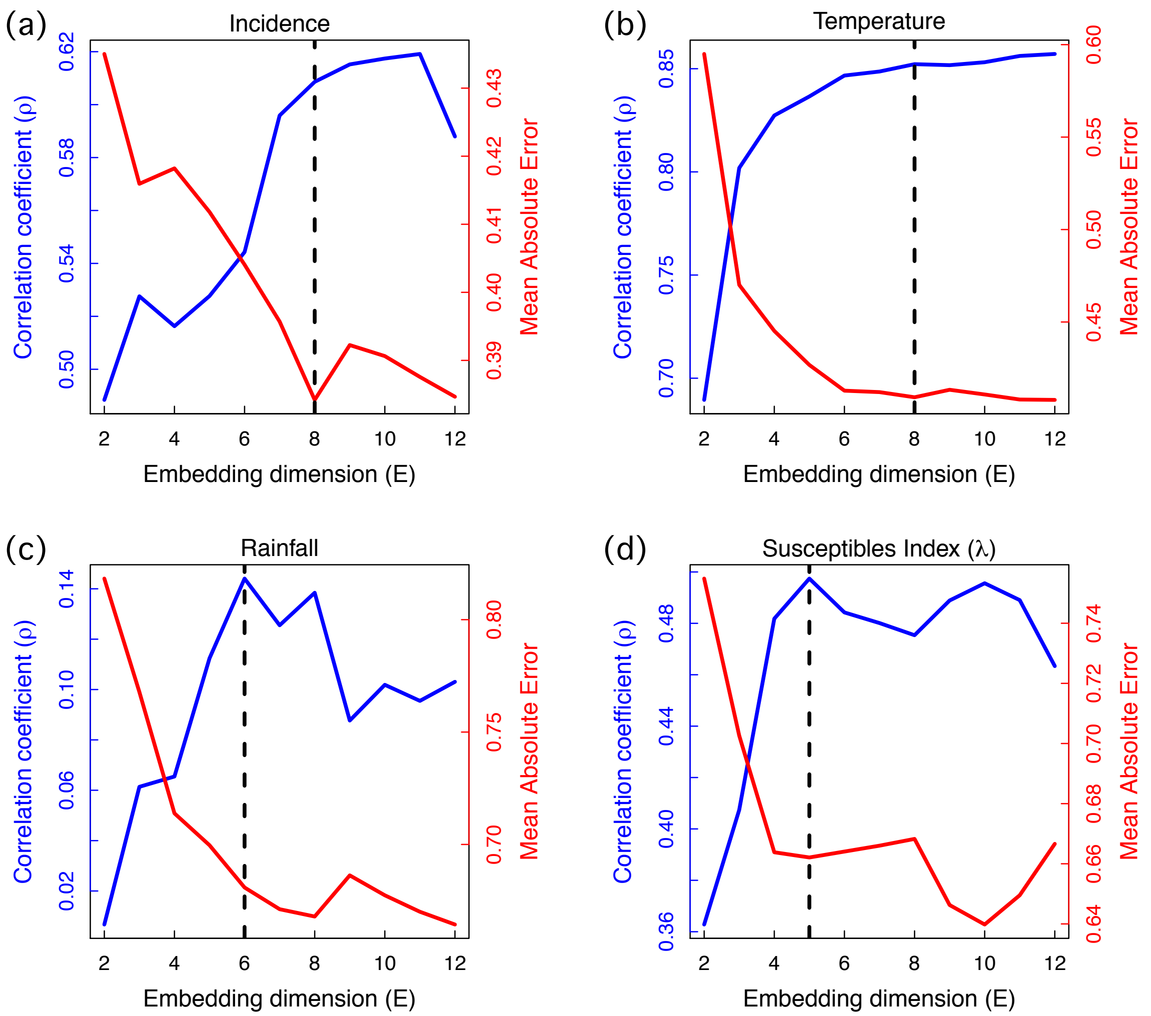


**Figure S3. Optimal embedding dimension (*E*).** For each univariate SSR, we calculated the correlation coefficient $\rho$ (blue lines) and mean absolute error (MAE; red lines) of predicted states (using simplex projection) versus observed states against different values of embedding dimension, $E$. The optimal embedding dimensions for incidence (a), temperature (b), rainfall (c) and the proxy for the susceptible population size (d) obtained here for further analyses were 8, 8, 6 and 5, respectively (dashed lines), i.e., approximately where $\rho$ peaks and MAE troughs.

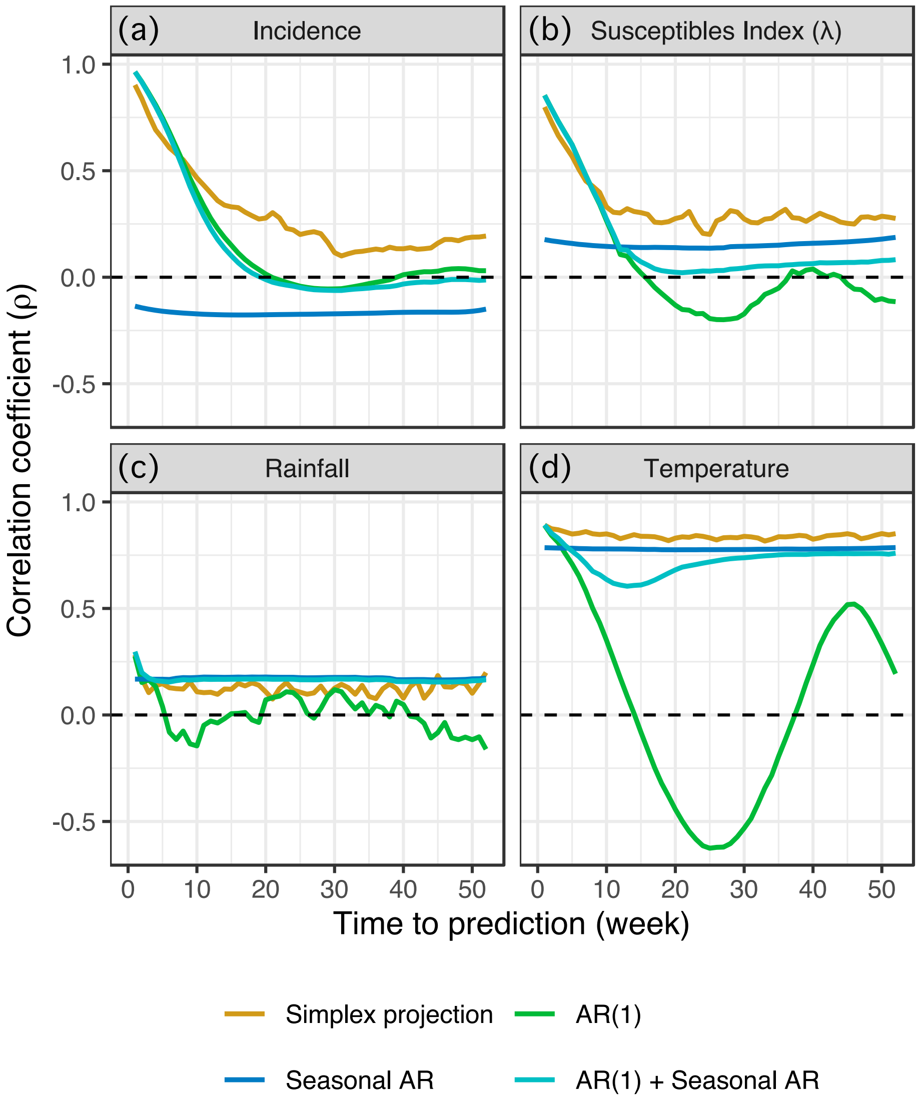


**Figure S4. Evidence of low-dimensional deterministic dynamics.** Shown is the correlation coefficient $\rho$ of predicted states against observed states against time to prediction using simplex projection (red), simple autoregressive model: $y(t) = \alpha y(t-1) + \epsilon(t)$ (green), seasonal autoregressive model: $y(t) = \beta y(t-52) + \epsilon(t)$ (cyan), and combined simple and seasonal autoregressive model: $y(t) = \alpha y(t-1) + \beta y(t-52) + \epsilon(t)$ (purple) for each variable state-space reconstruction: dengue incidence (a), temperature (b), rainfall (c), and the proxy for the susceptible population size (d). When time to prediction exceeds ~7 weeks, the system for incidence is forecastable (using simplex projection on an attractor) in a manner that exceeds the skill of an autoregressive model (a). This indicates that there is a signal of deterministic dynamics for incidence, since the prediction skill that accounts for attractor dynamics is above that of simple or seasonal autocorrelation. The presence of attractor geometry allows for better predictions using attractor-based methods (i.e., simplex projection)—an indication that an attractor (as a result of underlying deterministic dynamics) exists for incidence (a), susceptibles index (b), and temperature (d). By contrast, rainfall is a stochastic variable (c). However, EDM allows the use of stochastic drivers (e.g., rainfall) if an attractor exists for the response variable such as dengue incidence (a) (Munch *et al.* 2020).

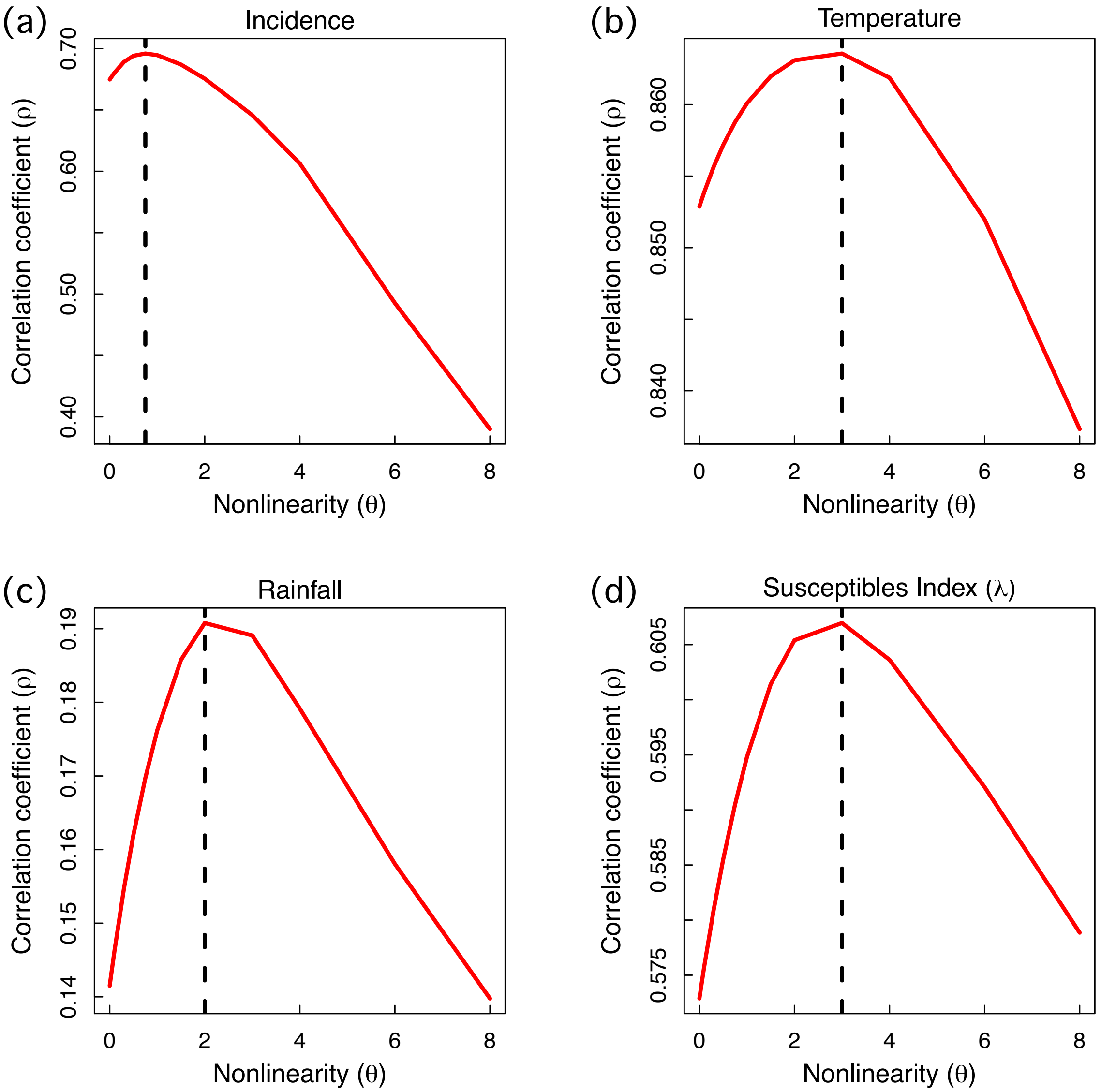


**Figure S5. Evidence of nonlinear dynamics.** The S-map test for nonlinearity confirms that all variables have nonlinear state dependence. Here we show the predictive skill ($\rho$) against the localization parameter $\theta$ (i.e., a scalar of the closeness between a predictor point and the point to be predicted on an attractor used in multivariable regression) for dengue incidence (a), temperature (b), rainfall (c), and proxy for susceptible population size (d). If the optimal $\theta > 0$ for a univariate state-space reconstruction of a variable, then local points around the point to be predicted are more important predictors (should have more weight) than other points further away on the attractor. This implies that the system is state-dependent and the variable is a part of a nonlinear dynamic system since predictor point location on the attractor affects prediction skill. Here, all variables have maxima at $\theta > 0$ (dashed line), which confirms that all variables have a nonlinear state dependence and thus motivates the use of EDM. If $\theta = 0$ then all points are weighted equally, and the system is linear.

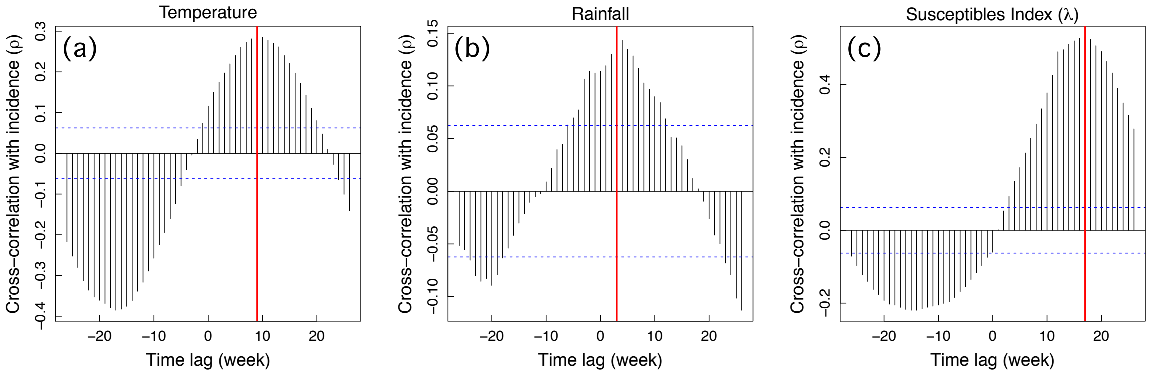


**Figure S6. Cross-correlations of drivers with dengue incidence.** Based on cross-correlation with dengue incidence, temperature has the optimal correlation at a 9-week lag (a; red line), rainfall at a 3-week lag (b; red line), and susceptibles index at a 17-week lag (c; red line). However, the susceptibles index time series were already shifted by 12 weeks due to the 12-week moving window when computing the index, so the actual seasonal lag is 17–12 = 5 weeks (Figure 2). Further, since the accumulation of rainfall over a longer period of time has biological significance as a potential driver of mosquito breeding habitat, we used the average rainfall of 3–9 weeks prior to incidence (7-week window; Figure 2). The cross-correlation was computed as $CCF(\tau) = \rho(Y(t),\ X(t+\tau))$ where $Y(t)$ is dengue incidence and $X(t+\tau)$ represents a driver (i.e., temperature, rainfall, or susceptibles index), and $\tau$ represents a time lag. A positive lag indicates that a driver is correlated with future dengue cases. The blue dashed lines represent the values beyond which the correlations are significantly different from zero with a 95% confidence interval.

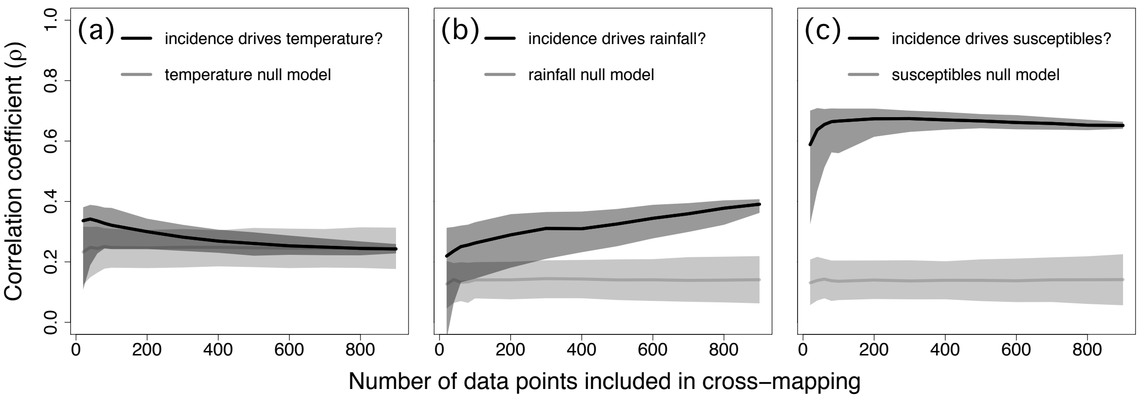


**Figure S7. Convergent cross-mapping correctly indicates that dengue incidence is not driving climate.** Cross-mapping showed that dengue incidence is not a driver of temperature (a; black), or rainfall (b; black) when testing the nonsensical direction of causality. However, incidence is a driver of the susceptibles index (c; black), since it makes sense that dengue incidence would affect the future susceptible population size. The cross-mapping skill (i.e., correlation coefficient $\rho$ between predicted driver values using the univariate state-space reconstruction of the response variable, and the observed driver values) did not converge (i.e., gradually increase until reaching a flat asymptote) as the number of time series data points increased. Lack of convergence indicates no causal relationship. The dark shaded regions represent the 0.025 and 0.975 quantiles of bootstrapped time series segments. The light grey shaded regions represent the 0.025 and 0.975 quantiles of the seasonal null distributions obtained from 500 runs of randomized time series with conserved seasonal trends (Deyle *et al.* 2016a). The grey line represents the median of the null distribution. Since there is no convergence (no causal forcing of dengue incidence on the other variables), the null models serve no purpose here, other than for comparing these results with Figure 3.

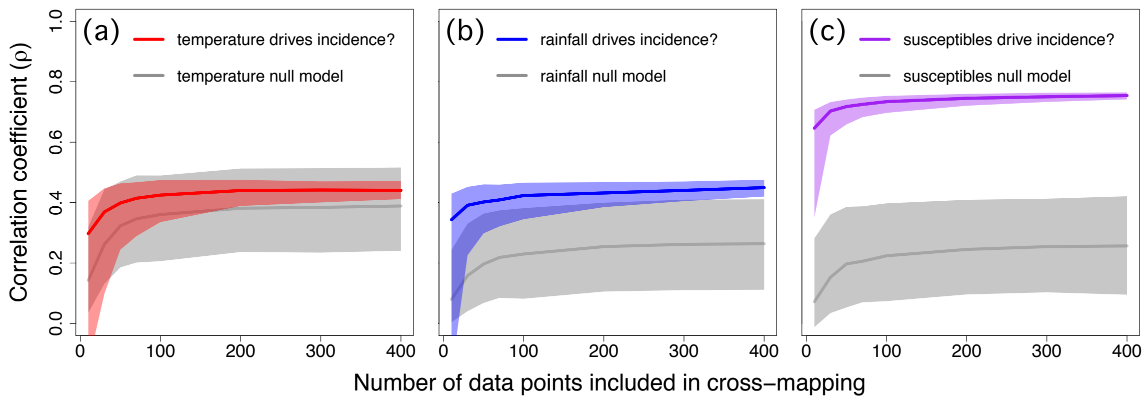


**Figure S8. Climate and susceptibles index drive dengue incidence (results using Ebisuzaki null model).** Cross-mapping between dengue incidence and temperature (a; red), rainfall (b; blue), or susceptibles index (c; purple) display significant (Kendall's $\tau > 0$; $P < 0.01$) convergence in cross-mapping skill (i.e., $\rho$ increases and reaches a flat asymptote) as the number of time series data points increases (sign of causality). Red, blue and purple shaded regions represent the 0.025 and 0.975 quantiles of bootstrapped time series segments. Grey shaded regions represent the 0.025 and 0.975 quantiles of the null distributions obtained from 500 runs of randomized time series with conserved cyclic trends (Ebisuzaki 1997). Solid lines represent medians of distributions. Rainfall and susceptibles index showed significant forcing above and beyond seasonal signal (K-S $P < 0.001$), because cross-mapping of the true time series (blue and purple) are distinguishable from their respective null models (grey).

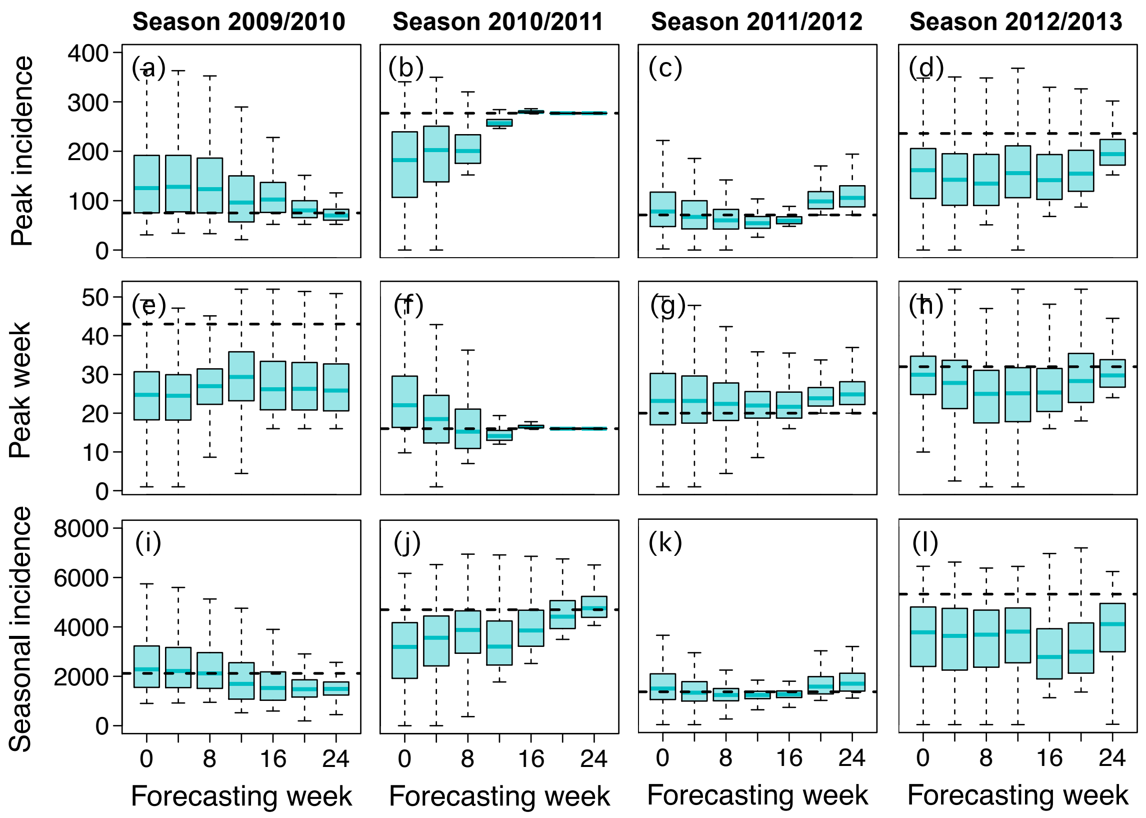


**Figure S9. Boxplots of forecasting target results using our forecasting model as specified in the dengue forecasting challenge for the testing period** (Johansson *et al.* 2019)**.** Forecasts were made at weeks 0, 4, …, 24 in all the testing seasons (2008/2009–2012/2013) for each target: peak incidence per season (a–d), peak week per season (e–h), and total number of cases per season (i–l). Each observed target value is indicated with a dashed horizontal line.

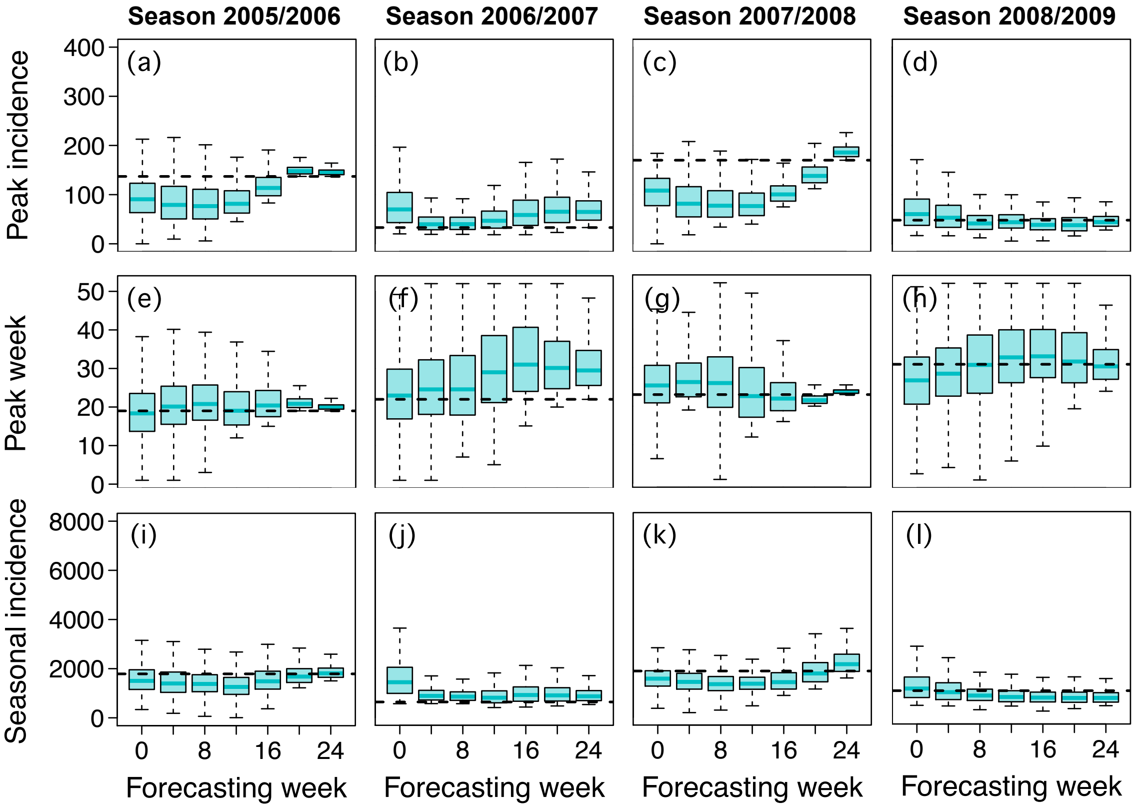


**Figure S10. Boxplots of forecasting target results using our forecasting model as specified in the dengue forecasting challenge for the training period** (Johansson *et al.* 2019)**.** Forecasts were made at weeks 0, 4, …, 24 in all the training seasons (2005/2006–2008/2009) for each target: peak incidence per season (a–d), peak week per season (e–h), and total number of cases per season (i–l). Each observed target value is indicated with a dashed horizontal line.

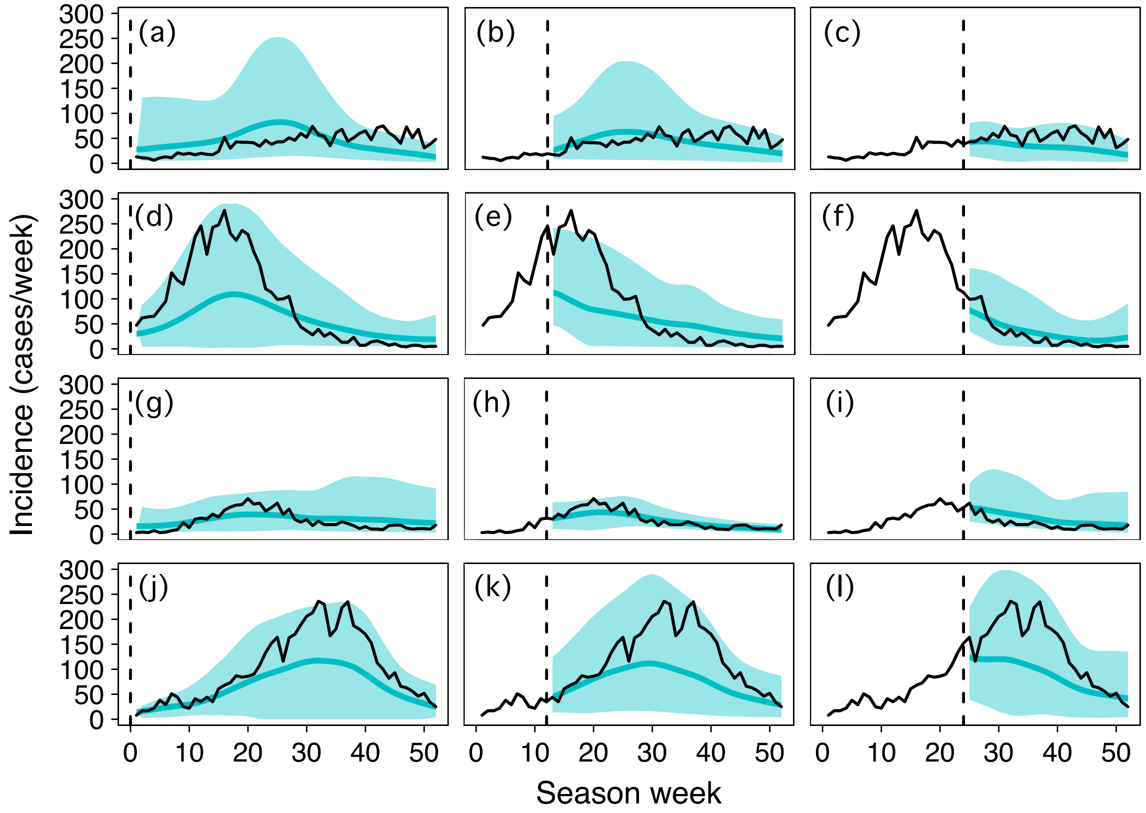


**Figure S11. Seasonal forecasts of our forecasting model during testing seasons (2009/2010–2012/2013).** Forecasts were made at weeks 0, 4, …, 24 in all the testing seasons and are shown in turquoise (solid lines represent the mean; shaded regions represent 95% confidence intervals) and observed incidence in black. Here we show the forecasts made at week 0 (a, d, g, j), week 12 (b, e, h, k), and week 24 (c, f, i, l) for seasons 2009/2010 (a, b, c), 2010/2011 (d, e, f), 2011/2012 (g, h, i), and 2012/2013 (j, k, l). Dashed vertical lines represent the weeks at which forecasts were made (i.e., 0, 12, or 24).

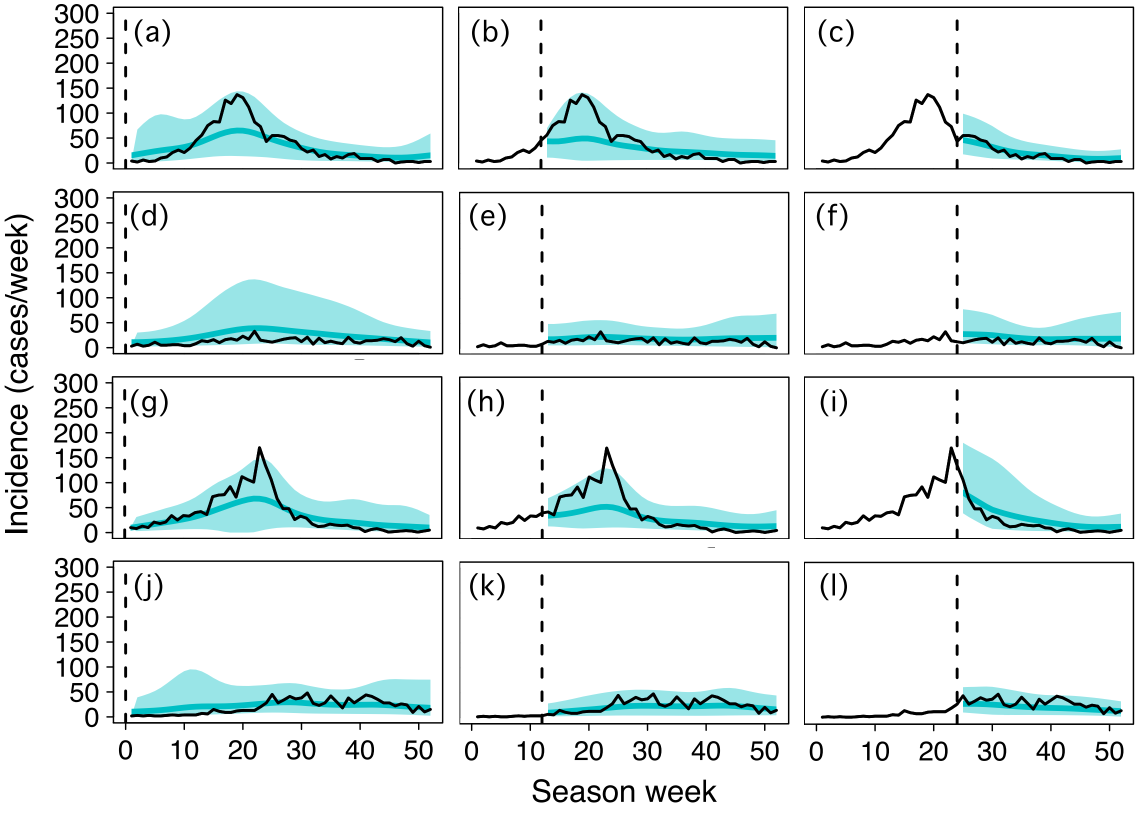


**Figure S12. Seasonal forecasts of our forecasting model during training seasons (2005/2006–2008/2009).** Forecasts were made at weeks 0, 4, …, 24 in all the training seasons and are shown in turquoise (solid lines represent the mean; shaded regions represent 95% confidence intervals) and observed incidence in black. Here we show the forecasts made at week 0 (a, d, g, j), week 12 (b, e, h, k), and week 24 (c, f, i, l) for seasons 2009/2010 (a, b, c), 2010/2011 (d, e, f), 2011/2012 (g, h, i), and 2012/2013 (j, k, l). Dashed vertical lines represent the weeks at which forecasts were made (i.e.,  0, 12, or 24).

### SUPPLEMENTARY TABLES

| Study | Peak Incidence | Peak Week | Seasonal Incidence | All targets (average) |
|---|---|---|---|---|
| Our approach | −1.02 | −3.11 | −1.26 | −1.79 |
| A | −4.62 | −6.02 | −4.79 | −5.14 |
| B | −2.57 | −4.24 | −2.04 | −2.95 |
| C | −2.12 | −4.14 | −2.13 | −2.80 |
| D | −2.62 | −6.45 | −5.07 | −4.71 |
| E | −1.43 | **−3.70** | −2.81 | −2.65 |
| F | −2.99 | −3.98 | −3.98 | −3.65 |
| G | **−1.23** | −4.88 | −1.992 | −2.70 |
| H | −2.60 | −5.37 | −2.64 | −3.54 |
| I | −2.95 | −6.03 | −3.32 | −4.10 |
| J | −1.74 | −4.20 | **−1.986** | **−2.64** |
| K | −4.49 | −6.17 | −4.38 | −5.01 |
| L | −3.30 | −5.45 | −4.19 | −4.31 |
| M | −5.17 | −5.75 | −7.66 | −6.19 |
| N | −4.98 | −4.06 | −2.18 | −3.74 |
| O | −2.66 | −3.90 | −2.84 | −3.13 |
| P | −2.63 | −5.36 | −4.55 | −4.18 |
| Null | −2.40 | −3.95 | −2.40 | −2.92 |
| Baseline | −1.43 | **−3.47** | −2.15 | **−2.35** |
| Ensemble (A–P) | −1.68 | −3.60 | −2.13 | −2.47 |

**Table S1. Comparison of our approach with other approaches in the dengue forecasting challenge for the testing period** (Johansson *et al.* 2019)**, where score values closer to zero indicate higher forecasting skill.** Average logarithmic forecasting scores for weeks 0, 4, …, 24 in all the testing seasons (2008/2009–2012/2013) for each target: peak incidence per season, peak week per season, and total number of cases per season, as reported from the forecasting challenge: Table S1 (Johansson *et al.* 2019). A score of zero indicates perfect forecasting (i.e., predictions equal the observations). The lowest (best) score reported in the forecasting challenge for each target is shown in bold (if the baseline outperformed all teams, then both the top team and the baseline scores are shown in bold). Scores from our study (highlighted in grey) outperformed all the other approaches reported in the forecasting challenge, as well as the ensemble (Johansson *et al.* 2019). A–P represent different models from teams that participated in the challenge, the null model assigned equal probably for every possible outcome, the baseline was a statistical time series model (a seasonal autoregressive integrated moving average [SARIMA] model), and the ensemble model output averages of the probabilities of all team model forecast and the baseline model forecasts.

| Study | Peak Incidence | Peak Week | Seasonal Incidence | All targets (average) |
|---|---|---|---|---|
| Our approach | −1.14 | −2.73 | −0.66 | −1.51 |
| A | −4.74 | −6.36 | −5.04 | −5.38 |
| B | −1.43 | −5.11 | −1.17 | −2.57 |
| C | −1.22 | −3.01 | −2.09 | −2.11 |
| D | −1.44 | −4.27 | −4.83 | −3.51 |
| E | −1.28 | −3.28 | −1.91 | −2.16 |
| F | −1.06 | −5.31 | −1.47 | −2.61 |
| G | −1.10 | −9.48 | −0.99 | −3.86 |
| H | −1.81 | −3.08 | −0.93 | −1.94 |
| I | −5.20 | −4.12 | −2.12 | −3.81 |
| J | −1.36 | −2.86 | −1.33 | −1.85 |
| K | −2.61 | −6.91 | −2.18 | −3.90 |
| L | −2.21 | −4.15 | −2.60 | −2.99 |
| M | **−0.94** | −3.54 | **−0.74** | −1.74 |
| N | −4.41 | −3.23 | −1.25 | −2.96 |
| O | −1.31 | **−2.74** | −0.91 | **−1.65** |
| P | −2.21 | −3.11 | −1.81 | −2.38 |
| Null | −2.40 | −3.95 | −2.40 | −2.92 |
| Baseline | −1.41 | −2.89 | −0.85 | −1.72 |
| Ensemble (A–P) | −1.24 | −2.93 | −0.97 | −1.71 |

**Table S2. Comparison of our approach with other approaches in the dengue forecasting challenge for the training period** (Johansson *et al.* 2019)**, where score values closer to zero indicate higher forecasting skill.** Average logarithmic forecasting scores for weeks 0, 4, …, 24 in all the training seasons (2005/2006–2008/2009) for each target: peak incidence per season, peak week per season, and total number of cases per season, as reported from the forecasting challenge: Table S2 (Johansson *et al.* 2019). A score of zero indicates perfect forecasting (i.e., predictions equal the observations). The lowest (best) score reported in the forecasting challenge for each target is shown in bold (if the baseline outperformed all teams, then both the top team and the baseline scores are shown in bold). Scores from our study are highlighted in grey. A–P represent different models from teams that participated in the challenge, the null model assigned equal probably for every possible outcome, the baseline was a statistical time series model (a seasonal autoregressive integrated moving average [SARIMA] model), and the ensemble model output averages of the probabilities of all team model forecast and the baseline model forecasts.
